# Biophysics and population size constrains speciation in an evolutionary model of developmental system drift

**DOI:** 10.1101/123265

**Authors:** Bhavin S. Khatri, Richard A. Goldstein

## Abstract

Developmental system drift is a likely mechanism for the origin of hybrid incompatibilities between closely related species. We examine here the detailed mechanistic basis of hybrid incompatibilities for a genotype-phenotype map for developmental system drift under stabilising selection, where the organismal phenotype is conserved, but the underlying molecular phenotypes and genotype can drift. This leads to number of emergent phenomenon not obtainable by modelling genotype or phenotype alone. Our results show that: 1) speciation is more rapid at smaller population sizes with a characteristic, Orr-like, power law, but at large population sizes slow, characterised by a sub-diffusive growth law; 2) the molecular phenotypes under weakest selection contribute to the earliest incompatibilities; and 3) pairwise incompatibilities dominate over higher order, contrary to previous predictions that the latter should dominate. Our results indicate that biophysics and population size provide a much stronger constraint to speciation than suggested by previous models.

## INTRODUCTION

The detailed genetic mechanisms by which non-interbreeding species diverge is still poorly understood. Darwin, inspired by John Herschel, called it that “mystery of mysteries” [16]; he struggled to understand how natural selection could give rise to hybrid inviability or infertility between populations without producing such incompatibilities within the populations. A solution to this problem was conceived independently by Dobzhansky, Muller and Bateson in which cross-mating would combine alleles at different loci that are incompatible due to epistatic interactions (Dobzhansky Muller incompatibilities, DMI) [5, 17, 58]. Consider a common ancestor with alleles ab across two loci, which after a period of allopatric divergence give rise to two lineages which have fixed genotypes *Ab* and *aB*, respectively Interbreeding between these two populations would result in the heterozygotic hybrid genotype *Aa*|*Bb*, combining the potentially incompatible *A* and *B*, a combination that could not arise in either population mating separately.

Assuming that any combination of alleles that have not been “tested” by the process of evolution represents a potential incompatibility Orr predicted that the number of incompatibilities between pairs of alleles in a sufficiently large genome would increase with the number of substitutions separating the two lineages (*K*) as *K*(*K* − 1) ~ *K*^2^ [64]. Similarly the number of untested combinations involving *n* loci would increase as ~ *K^n^*, suggesting that, with evolutionary time, potential incompatibilities would become increasingly dominated by more complex epistatic interactions [64]. This would occur, firstly, because there are a larger number of combinations, and secondly, because there are more ways for separate lineages to evolve around incompatible genotypes when there is a larger number of loci. It is unclear, however, how the simplistic assumptions of this model fare with increased biological realism. Not all possible untested hybrids are equally likely to result in real incompatibilities. Selection acting on each separate lineage affects the substitutions that occur and their likely contributions to reproductive isolation. In particular, evolutionary constraints have a strong effect on the development of more complex DMIs, making it uncertain whether their role is as important as suggested by Orr’s combinatorial argument. This highlights the need for considering more realistic models that better capture the salient aspects of the underlying biology, whilst remaining sufficiently simple for tractable evolutionary modelling and simulation.

In recent years a form of epistasis has been described in a number of organisms whereby closely related species have similar organismal phenotypes but are produced by very different developmental mechanisms [56, 79]. This cryptic “developmental system drift” [33, 76] could be an important source of hybrid incompatibilities that cause reproductive isolation [38, 39]. Developmental system drift is an example of a more general characteristic of biological systems where many genotypes can correspond to the same phenotype; this redundancy of the mapping from genotype to phenotype results in a number of non-trivial behaviours which do not arise on fitness landscapes which consider evolution of phenotypes or genotypes independently [7, 23, 29–32, 35, 44, 52, 55, 59, 69]. The degree of redundancy can be represented as the “sequence entropy”, corresponding to the log of the number of genotypes corresponding to a given phenotype, in analogy to the similar expression in statistical mechanics [30, 42–44].

To explore the role of developmental system drift on speciation, we examine the growth of Dobzhansky-Muller incompatibilities using a simple genotype-phenotype map that models the development of spatial patterning of gene expression. The model, introduced by [44] allows for cryptic genetic variation and changes in molecular phenotypes while maintaining organismal phenotype under stabilising selection. In addition, we introduce a novel computational method to decompose hybrid DMIs so we can examine the behaviour of the fundamental pair-wise and higher order incompatibilities. We show that including biologically rele vant elements gives rise to a number of novel phenomenon that could not arise with models based only on the fitness of genotypes or phenotypes. Our results show that small populations develop hybrid incompatibilities more quickly, due to the pressure of sequence entropy in small populations meaning the common ancestor harbours on average a larger drift load. For large populations, we find hybrid incompatibilities arise more slowly, with a growth law characteristic of a sub-diffusion of the hybrid binding energies, indicative of kinetic traps in the molecular substitution process due to roughness to the fitness landscape [44]. Strikingly, we find that for moderate population sizes it is the molecular phenotypes under weakest selection that give rise to earliest incompatibilities, since in the common ancestor they are more likely to be already maladapted. Finally, we find that unlike Orr’s prediction that complex DMIs should be abundant, pair-wise interactions between loci dominate the growth of DMIs, showing that biophysics provides a stronger constraint than pure combinatorics.

## A SIMPLE GENOTYPE-PHENOTYPE MAP TO MODEL DEVELOPMENTAL SYSTEM DRIFT

The genotype-phenotype map we use is a modification of the one described in [44]. The evolutionary task set for the gene regulation module is to turn an exponentially decaying morphogen gradient (*M*) across a field of cells in an embryo into a sharp step function profile of a downstream transcription factor *T* with its transition at the mid-point of the embryo, as shown in Fig.1. This is accomplished by having the morphogen and an RNA Polymerase *R* bind to two adjacent non-overlapping binding sites in the cis-regulatory region (*C*) region of the transcription factor, the promoter *P*, and a single binding site *B* adjacent to it; transcription occurs whenever the polymerase binds to the promoter, although both proteins can bind to both binding sites dependent on their binding affinities. Binding to the regulatory region is cooperative due to stabilising interactions between the two proteins when bound at the two adjacent sites. The sequences of *M* and *R* at the DNA binding sites are represented by binary strings of length *ℓ_pd_* = 10. The corresponding DNA binding sites *B* and *P* are also represented by binary strings of the same length. Interactions between a pair of proteins are similarly represented by binary strings of length *ℓ_pp_* = 5. We assume an exponential morphogen concentration profile [*M*](*x,α*), as a function of the position of embryonic cells, *x*; the decay rate of the morphogen *α* is represented as a continuous variable, with a relative probability of mutation corresponding to an effective string of length *ℓ_α_* = 10 bases. This results in a genome ***G***, of total length *ℓ_**G**_* = 60. Protein-DNA and protein-protein binding strengths are determined by the number of mismatches between corresponding strings on the two interacting molecules, where for protein-DNA binding the cost of a mismatch is *ϵ_pd_* = 2*k_B_T* and for protein-protein interactions *ϵ_pp_* = 1 *k_B_T*, where *k_B_T* is Boltzmann’s constant multiplied by room temperature (298K). We assume that there is a fixed concentration [*R*] of polymerase, in each cell. We then follow [71] and assume that the concentration of the transcription factor in a cell at position *x*([*T*](*x*)) is simply proportional to the probability of the polymerase being bound to the promoter. The fitness contribution *F* of the overall patterning phenotype ranges from 0 to *κ_F_* depending on how well expression of the transcription factor is confined to the anterior half of the embryo, as shown in Fig. 1 (bottom left), where *κ_F_* is a measure of the relative contribution of this trait to the fitness of the organism. We define a population-scaled fitness contribution 2*N_e_κ_F_*, where *N_e_* is the effective population size; for 2*N_e_κ_F_* < 1 the effects of selection are weak, and are conversely strong when 2*Nκ_F_* > 1. We also assume that there is a boundary at *F* = *F*\*, below which the organism is unviable. We simulate evolution as continuous time Markov process. After evolving a single population for a given number of generations, we form two replicates of the population that evolve independently, representing the process of allopatric speciation. At various time points following this imposed isolation, we consider the fitness and viability of various outcrossings between the two populations. A DMI occurs when the fitness contribution of a particular hybrid drops below *F*\*.

**FIG. 1.**
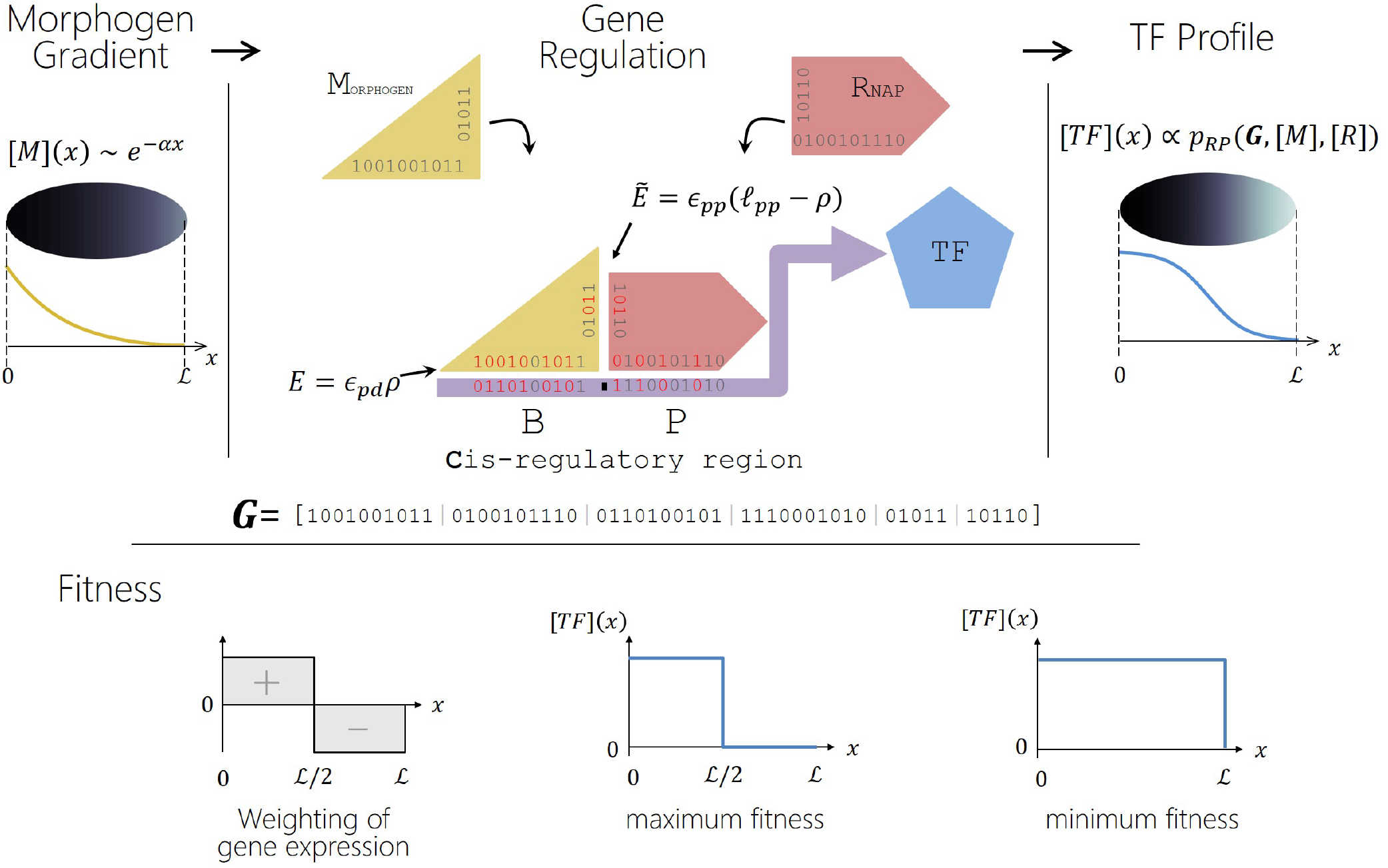
An overview of the genotype-phenotype map. The gene regulatory module has as input a morphogen concentration [*M*](*x*) that varies approximately exponentially across a 1-dimensional embryo of length *L*, and outputs a transcription factor [*T*](*x*). Spatial gene regulation of *T* is achieved via a simple Hamming model of protein DNA binding as shown. Fitness is determined by weighting any gene expression in the anterior half of the embryo as positive, and any in the posterior half as negative (as shown bottom left). We show idealised expression profiles that give maximum fitness when gene expression is confined only to the anterior half (bottom middle), and minimum fitness when there is no discrimination between anterior and posterior (bottom right); these translate to log-fitnesses of *F* = 0 and *F* = −∞, respectively (see Methods).

## RESULTS

### Evolutionary properties of genotype-phenotype map on each lineage

The properties of a similar genotype-phenotype map have been previously explored [44]. An important property of this genotype-phenotype map is that only a single mechanism of patterning is found, in which the polymerase (*R*) binds with intermediate affinity to the promoter (*P*) but with high affinity to the morphogen (*M*), while the morphogen binds to the morphogen binding site (*B*) only above a critical morphogen concentration. This results in a spatial switch once the morphogen falls below this concentration; evolution then fine tunes the relationship between the protein-DNA binding energies, the protein-protein binding energy and the steepness of the morphogen gradient *α* to turn off transcription at the mid-point of the embryo. Despite a single global solution there are many different combinations of the protein-DNA and protein-protein binding energies and *α* that give good patterning, and each of these correspond to many possible genotypes (***G***). Of the different possible binding energies, we find that *E_MB_*, *E_RP_*, *Ẽ_RM_* are under strong selection, whilst the other possible binding energies are essentially neutral with weak selective effects. At large population sizes it is found that the evolutionary dynamics exhibits what is known as quenched-disorder in statistical physics, where energy phenotypes that are less constrained take different *random* values between independent evolutionary runs with no further substitutions; this indicates an underlying roughness to the fitness landscape and that these non-important trait values in a local optimum [44].

A key property determining the rate at which incompatibilities arise is the distribution of common ancestor phenotypes as a function of the population-scaled fitness contribution 2*N_e_κ_F_*, as shown in Fig. 2. For a given value of *κ_F_*, we see that for large population sizes (2*Nκ_F_* ≫ 10) the distribution is what we would expect from conventional evolutionary theory on a fitness landscape with a fitness maximum. In contrast, as the population size is decreased, we find the distribution shifts to lower fitness values to the point when selection is weak (2*Nκ_F_* ≤ 1) the distribution is poised at the inviability boundary. This effect arises due to genetic drift at low population sizes pushing populations towards marginally fit phenotypes that correspond to the largest number of genotypes, that is, with the largest sequence entropy.

**FIG. 2.**
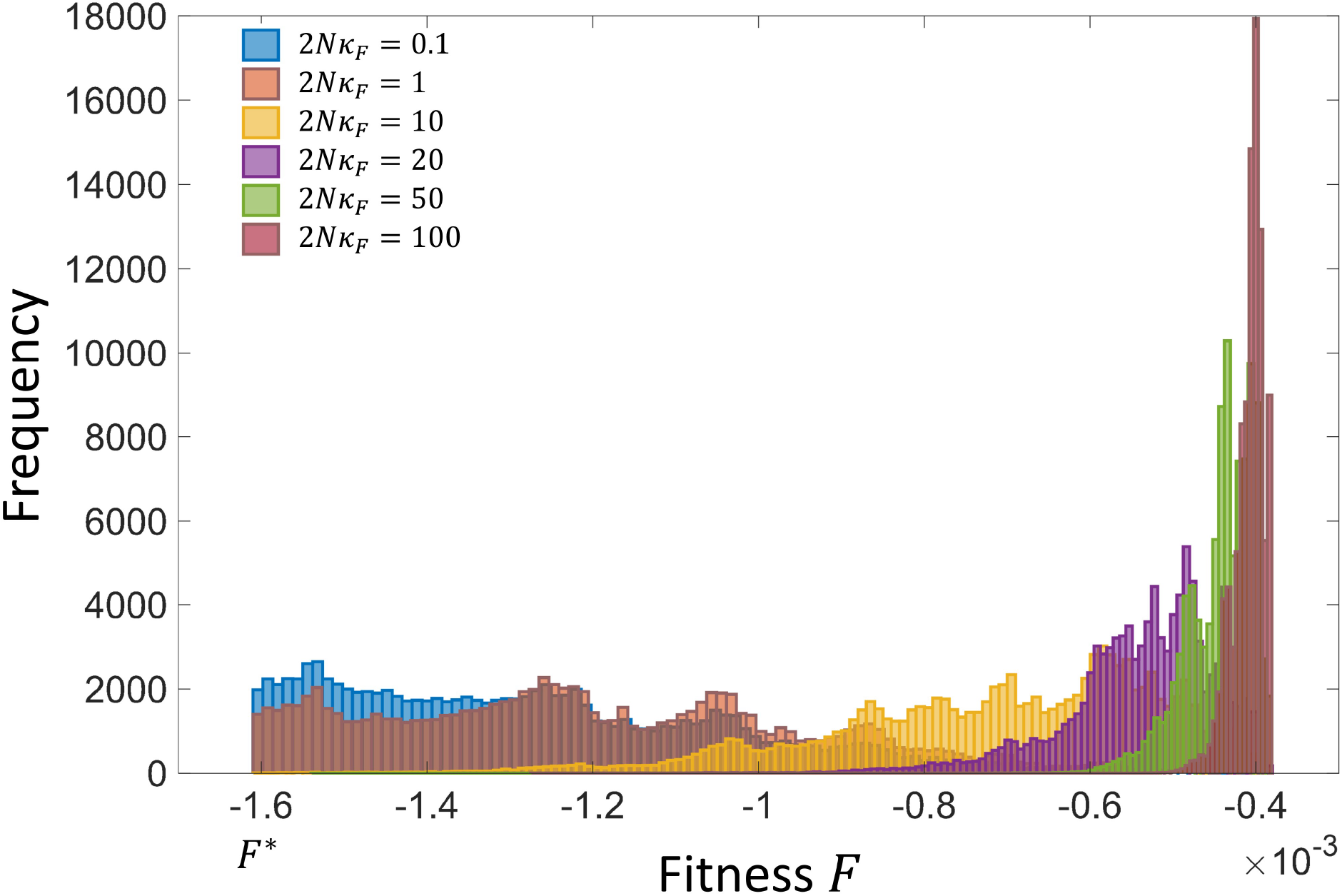
Histogram of the fitness of populations over single long runs as a function of the population scaled fitness contribution 2*Nκ_F_*

### Decomposition of DMIs

Our genome is composed of 4 loci: 1) the *R* locus corresponding to the polymerase sequence, 2) the Morphogen (*M*) locus, 3) the *C* locus which corresponds to the sequences for the cis-regulatory region of the transcription factor and 4) the *α* locus, which is the morphogen gradient steepness *α*. Hybrids between the two lineages are constructed by independent reassortment of these loci assuming complete linkage within each locus and no linkage between them. We define a hybrid genotype by a 4 letter string where each letter corresponds to one of the loci defined above and takes one of two cases correspond to whether the allele is from the 1st line or 2nd line; for example, the hybrid rMCa corresponds to *R* locus having an allele from the 1st lineage, *M* locus with the allele from the 2nd lineage, the transcription factor (cis-regulatory) *C* locus from the 2nd lineage and α locus the allele from the 1st lineage. Note that the underlying sequence of each hybrid changes as different substitutions are accepted in each lineage; the notation only refers to alleles fixed at any point in time. We can represent all combinations of the four loci drawn from the two parents (RMCA, RMCa, RMcA, etc.) as points on a four-dimensional Boolean hypercube. In total there are 2^4^ − 2 = 14 hybrids.

In Fig.3, we plot a typical time series of how the fitness of two different hybrids (Rmca(a & b) and RMcA(c & d) changes over a divergence time *μt* separating a pair of lineages, for 2*Nκ_F_* = 1 (a & c) and 2*Nκ_F_* = 10 (b & d), where *μ* = *ℓ_**G**_μ*_0_ is the mutation rate for all base pairs in all loci. 2*Nκ_F_* > 1 indicates strong selection, whilst 2*Nκ_F_* ≤ 1 indicates weak selection where genetic drift dominates (For reference, in human populations it has been estimated that ≈ 20 − 30% of mutations are weakly selected [19, 36], compared to in *Drosophila* < 10% [36].). We see that the fitness of hybrids generally decreases in a stochastic fashion; when the log-fitness of a hybrid drops below the threshold *F*\* (indicated by the dashed line), a DMI arises as is indicated by a vertical log-fitness line (*F* = −∞) for that hybrid. As can be seen in Fig.3, at any given time a changing subset of the fourteen possible hybrids might be incompatible.

**FIG. 3.**
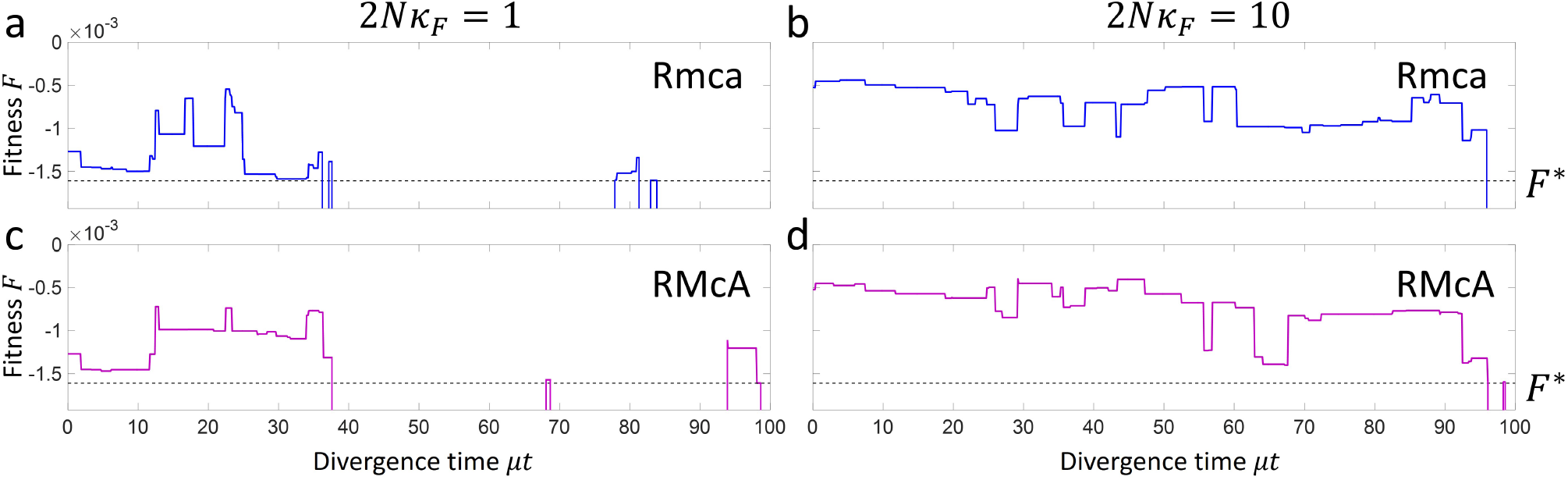
Plot of the times series of two hybrids Rmca(a & b) and RMcA(c & d) at population scaled fitness contributions of 2*Nκ_F_* = 1 (a & c) and 2*Nκ_F_* = 10 (b & d). Times when the fitness of a hybrid drop below the critical value *F*\* (dashed line) correspond to DMIs.

#### Total number and type of DMIs

To decompose the pattern of hybrid DMIs into fundamental pair-wise, 3-way and 4-way incompatibilities, we use a parsimonious method (Material & Methods) which finds the minimum total number of fundamental DMIs types that can explain the pattern of DMIs observed. As shown in Fig.4a, on a 4-dimensional Boolean hypercube, a pair-wise incompatibility is represented by a face of the cube (blue square), a 3-way incompatibility by a single edge (green line) and a 4-way incompatibility by a single point (red circle). This arises because, for example, as in Fig.4a, a 2-point DMI *I*_Ca_ means any genotype which features this sub-genotype Ca must, by definition, also be a DMI; the alleles on the remaining loci not involved in the DMI, can take any value and so form a 2*D* subspace, which is a whole face of the Boolean 4-cube. Similarly, a 3-point DMI, such as *I*_mcA_ as shown, constrains the sub-genotype of 3 loci to be incompatible and the remaining locus can take any value forming an edge (r → R in the example) of a 4*D* Boolean hypercube. A 4-point DMI must be a single point in a 4*D* Boolean hypercube (e.g. *I*_rmCA_ as shown). In Fig. 4b, we show an example where the pattern of hybrid incompatibilities, shown by red crosses, can be explained in three different ways, each with only 2 DMIs: 1) *I*_Ca_ + *I*_Rca_, 2) *I*_Ra_ + *I*_Ca_, 3) *I*_Ca_ + *I*_Ra_; two is the minimum number of DMIs, as there is no way to explain the pattern seen with a single pair-wise, 3-way or 4-way DMI. We assume each of these is, a priori, equally likely and so the total number of pair-wise incompatibilities is calculated as an average over the different ways we can explain the observed pattern of hybrid DMIs; in the example in Fig. 4b, we therefore have a total number of 2-way DMIs of *n*_2_ = 4/3 and 3-way DMIs *n*_3_ = 2/3, giving a total number of *n* = *n*_2_ + *n*_3_ = 2 DMIs, which is the minimum number of DMIs needed to explain the pattern of hybrid incompatibilities.

**FIG. 4.**
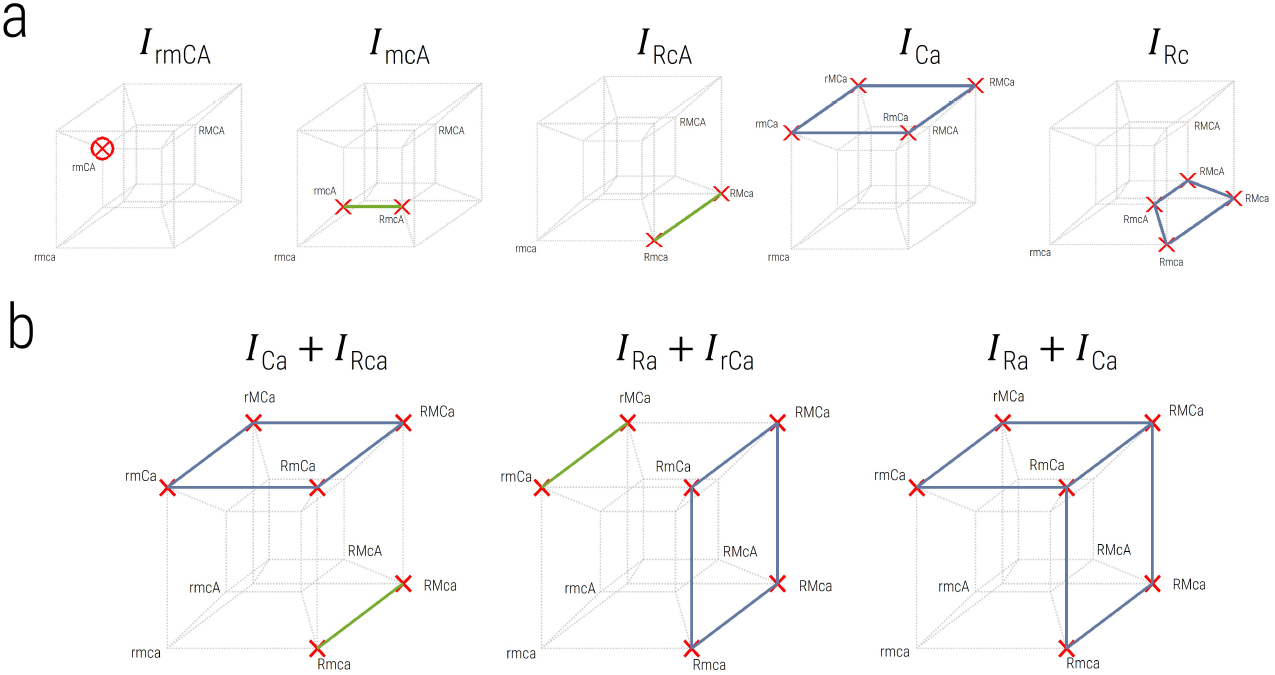
Decomposition of hybrid DMIs on a Boolean hyper 4-cube. Each point on the 4-cube represent each possible hybrid genotype across 4 loci, including the genotype of each parental lineage, where each red cross represents an incompatible hybrid genotype. As shown in a) the pattern of DMIs can be explained by different combinations of fundamental types of DMIs, where a blue square or face identify a subspace of genotypes that correspond to a 2-way DMI, a single green edge or line corresponds to a 3-way DMI and a red open circle corresponds to a single isolated 4-way DMI. b) A more complicated pattern of hybrid DMIs and their decomposition into fundamental types.

Using this method, we plot the total number of each type of DMI versus divergence time in Fig.5, where the panels correspond to different population scaled fitness contributions from 2*N_e_κ_F_* = 0.1 to 2*N_e_κ_F_* = 100. We see clearly that as the population size is decreased the rate at which incompatibilities arise increases. This effect results from the shift in distribution of fitness to poorer values for smaller populations in Fig. 2; this means that the common ancestor is more likely to be slightly maladapted (higher mutational load) for smaller populations, and so fewer substitutions are required before hybrid incompatibilities arise. This effect was also found for simple models of protein-DNA binding [42, 43], but arose as a result of direct selection for high binding affinity, which also corresponds to low sequence entropy as there are exponentially fewer ways of binding with no or only a few mismatches between protein and DNA. In this work there is only selection on the overall patterning phenotype, yet remarkably we find analogous behaviour due to the effect of sequence entropy of this phenotype.

**FIG. 5.**
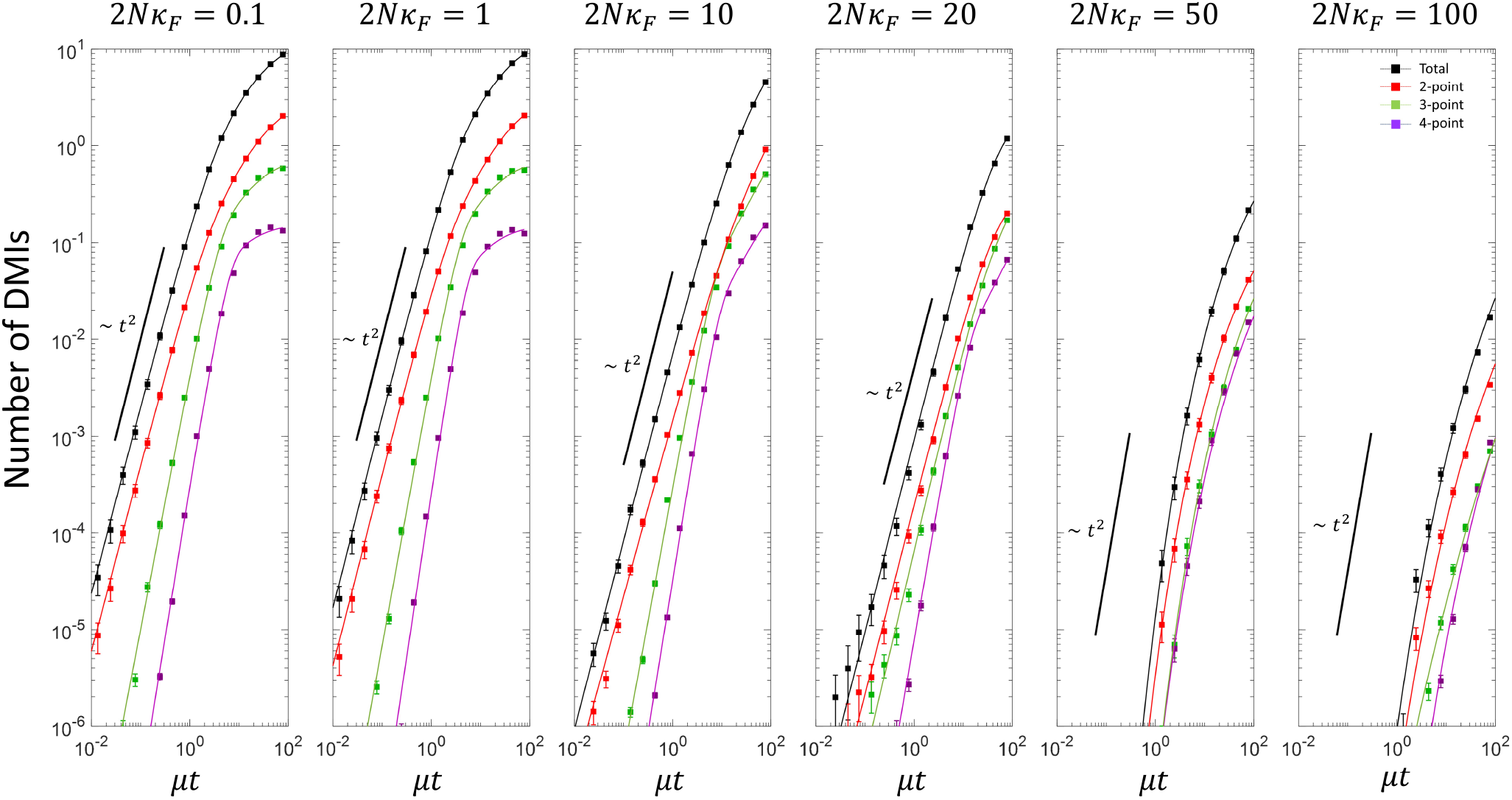
Plot of the total number of DMIs vs divergence time, together with their decomposition into the total number of 2-way, 3-way, 4-way DMIs, for various scaled populations sizes. For 2*Nκ_F_* ≤ 20 the solid lines correspond to fits of the simulation data to Eqn.1, while for 2*Nκ_F_* ≥ 50 correspond to fits to Eqn.2.

#### Pair-wise incompatibilities dominate reproductive isolation

As is clearly shown by Fig.5, pair-wise incompatibilities dominate for all population sizes, though at larger population sizes the difference is smaller, and particularly at shorter times. These results should be contrasted with the prediction of Orr that 2-way: 3-way: 4-way incompatibilities should arise in the ratio 12 : 24 : 14 (for *L* = 4 loci); so for example, there should be double the number of 3-way incompatibilities to pair-wise.

It is possible that this bias could arise by the development of a random set of incompatibilities that would preferentially decompose into sets of 2-way DMIs. In order to consider this possibility, we randomly assigned a set of incompatibilities and analysed them using our decomposition procedure. Contrary to the observed results,, we find that this decomposition results in the largest number of 4-way DMIs, followed by 3-way and then 2-way DMIs (Supporting Information). This is as expected as when the probability that a hybrid is incompatible is small, by random chance, we should expect to see isolated incompatible hybrid genotypes, which would be explained by 4-way DMIs; the fact that our results show that 2-way DMIs are dominant, even at early times, means that whole faces of the Boolean hyper-cube are found to incompatible, which is not likely to arise in a random model. Hence overall, these results show that contrary to Orr’s prediction that higher order DMIs should be easier to evolve, higher order DMIs evolve more slowly and are in smaller number compared to pair-wise DMIs.

#### Quantification of power law growth of DMIs for small populations

We find that for small population sizes 2-way, 3-way and 4-way DMIs all increase as a power law at small times, indicated by a straight line on a log-log plot, in agreement with [64, 66], who predicted that n-way DMIs should increase as ~ *t^n^*. To more quantitatively assess the exponent, we fit the data for 2*Nκ_F_* ≤ 20 using the phenomenological equation:

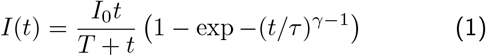

which has the asymptotic form of *I*(*t*) ~ *t^γ^* for *t* ≪ *τ* and *t* ≪ *T* and an opposite limit of *I*(*t* → ∞) = *I*_0_. We see that for the total number of DMIs and for 2-way, 3-way and 4-way DMIs, this form fits the data well at intermediate and small population sizes. We tabulate the power law exponent derived from these fits in Table I. We see that the total number of DMIs and 2-way DMIs have a power law exponent close to *γ* = 2, which is consistent with the Orr model. The 4-way DMIs have a larger exponent than the 3-way DMIs, which is larger than the 2-way DMIs, also in agreement with Orr’s model, although the values of the exponent are substantially lower than *n* for the 3-way and 4-way DMIs.

**TABLE I.** Table of values of the exponent γ characterising the power law of growth of DMIs at short times and small scaled population sizes.

| $2N\kappa_F$ | | 0.1 | 1 | 10 | 20 |
| --- | --- | --- | --- | --- | --- |
| Total | $\gamma$ | $1.94 \pm 0.05$ | $1.99 \pm 0.05$ | $1.95 \pm 0.05$ | $2.00 \pm 0.12$ |
| 2-way | | $1.93 \pm 0.05$ | $1.98 \pm 0.06$ | $1.84 \pm 0.06$ | $1.97 \pm 0.13$ |
| 3-way | | $2.66 \pm 0.13$ | $2.81 \pm 0.17$ | $2.76 \pm 0.24$ | $2.14 \pm 0.15$ |
| 4-way | | $3.17 \pm 0.19$ | $3.43 \pm 0.41$ | $3.14 \pm 0.24$ | $2.93 \pm 0.19$ |

#### Sub-diffusive growth of DMIs for large populations

For large population sizes there is no clear power law, which is again consistent with previous simulations [43] and also theoretical calculations [42]; when high fitness corresponds to high binding affinity, so that the common ancestor distribution is peaked away from the inviability boundary, large populations have a small drift load, meaning that DMIs arise when hybrid energy traits diffuse to the boundary. One such analytically tractable model was investigated in [42] and predicted that the number of DMIs is a complementary error function, which has an asymptotic form 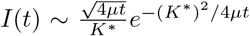, which due to the essential singularity as *t* → 0 has the property of negative curvature on a log-log plot. However, neither this form nor its multidimensional generalization fit the simulation data well (not shown). A functional form that *is* a good fit to the data at large populations sizes is

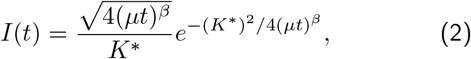

which arises when considering fractional Brownian processes with exponent *β* [8]; normal diffusion or Brownian motion arises when *β* =1, while *β* < 1 corresponds to subdiffusive behaviour, while *β* > 1 is superdiffusive. It is clear by examining the exponent *β* in Table II from fits of the data in Fig.5 at large population size (2*Nκ_F_* > 50) that the DMIs arise as a result of a sub-diffusive process, where 2-way, 3-way and 4-way DMIs have an exponent *β* ≈ 1/3 for 2*Nκ_F_* = 50 and *β* ≈ 1/4 for 2*Nκ_F_* = 100. The most likely mechanism that would give rise to sub-diffusive behaviour is a broad spectrum of times between substitutions; even though in the simulations the kinetic Monte Carlo scheme is based on a Poisson process for a given genotypic state ***G***, the distribution of rates could vary significantly as populations explore the fitness landscape. This would be consistent with the results in [44], which reveal the underlying fitness landscape of this spatial patterning genotype-phenotype map to be rough, which could lead to broad distribution of substitution rates in each lineage and effective sub-diffusive behaviour of the hybrids. Finally as expected the average number of substitutions needed at large population sizes is large, with values of *K*\* ranging from 6 to 9, and increases with increasing population size, as expected; it also increases very moderately with increasing *n*, which would be consistent with an increase in dimensionality of higher order DMIs. Interestingly, these values of *K*\* would indicate that the fraction of viable genotypes is very small, ~ 2^*K*\*^/2^*ℓ**G***^ ~ 10^−13^, where *ℓ_**G**_* = 60 is the number of effective binary sites in this genotype-phenotype map.

**TABLE II.** Table of values of the parameters characterising the sub-diffusive growth of DMIs for large scaled population sizes; *β* = 1 corresponds to normal diffusive motion, *β* < 1 to sub-diffusion and *β* > 1 super-diffusion, while *K*\* corresponds roughly to the number of substitutions required to reach the inviable region.

| $2N\kappa_F$ | | 50 | 100 |
| --- | --- | --- | --- |
| Total | $\beta$ | $0.47 \pm 0.03$ | $0.33 \pm 0.02$ |
| | $K^*$ | $6.58 \pm 0.12$ | $7.37 \pm 0.26$ |
| 2-way | $\beta$ | $0.32 \pm 0.01$ | $0.25 \pm 0.01$ |
| | $K^*$ | $6.71 \pm 0.07$ | $7.51 \pm 0.18$ |
| 3-way | $\beta$ | $0.33 \pm 0.01$ | $0.22 \pm 0.01$ |
| | $K^*$ | $7.53 \pm 0.10$ | $8.09 \pm 0.27$ |
| 4-way | $\beta$ | $0.31 \pm 0.02$ | $0.25 \pm 0.02$ |
| | $K^*$ | $7.50 \pm 0.28$ | $8.66 \pm 0.39$ |

### Molecular phenotypes under weakest selection dominate DMIs at larger population sizes

The DMI decomposition algorithm allows examination of the behaviour of different types of pair-wise, 3-way and 4-way DMIs. The different pair-wise DMIs can easily be identified with the different molecular binding energy phenotypes that combine together to form the overall organismal phenotype. The plot in Fig.6 shows how the number of DMIs grows for the three most dominant pair-wise DMIs *I*_rm_, *I*_rc_ and *I*_mc_ (pair-wise DMIs involving the *α* locus are orders of magnitude smaller - see Supporting Information), as a function of the scaled divergence time *ℓ*′*μt*, where *ℓ*′ = 5 for *I*_rm_ and *ℓ*′ = 10 for *I*_rc_ and *I*_mc_, cancelling how mutations are more likely to hit longer regions of the genome [43]. Strikingly, we find that contrary to what might seem intuitive, it is the molecular interactions under weakest selection that dominate the number of DMIs, particularly for larger population sizes. For 2*Nκ_F_* >= 50, we see at early times *I*_rc_ > *I*_rm_ > *I*_mc_, which is the same order of increasing selective constraint on each of these molecular interactions (Supporting Information). This has a simple explanation: at a sufficiently large population size such that none of the selective constraints on different molecular phenotypes is neutral, in the common ancestor the molecular interaction strengths under weakest selection will have the largest drift load and hence fewer substitutions are needed for incompatibilities to arise in hybrids. This is precisely as observed in simple models of transcription factor DNA binding, where the incompatibilities arise more quickly where binding is under weak selection for high affinity [43].

**FIG. 6.**
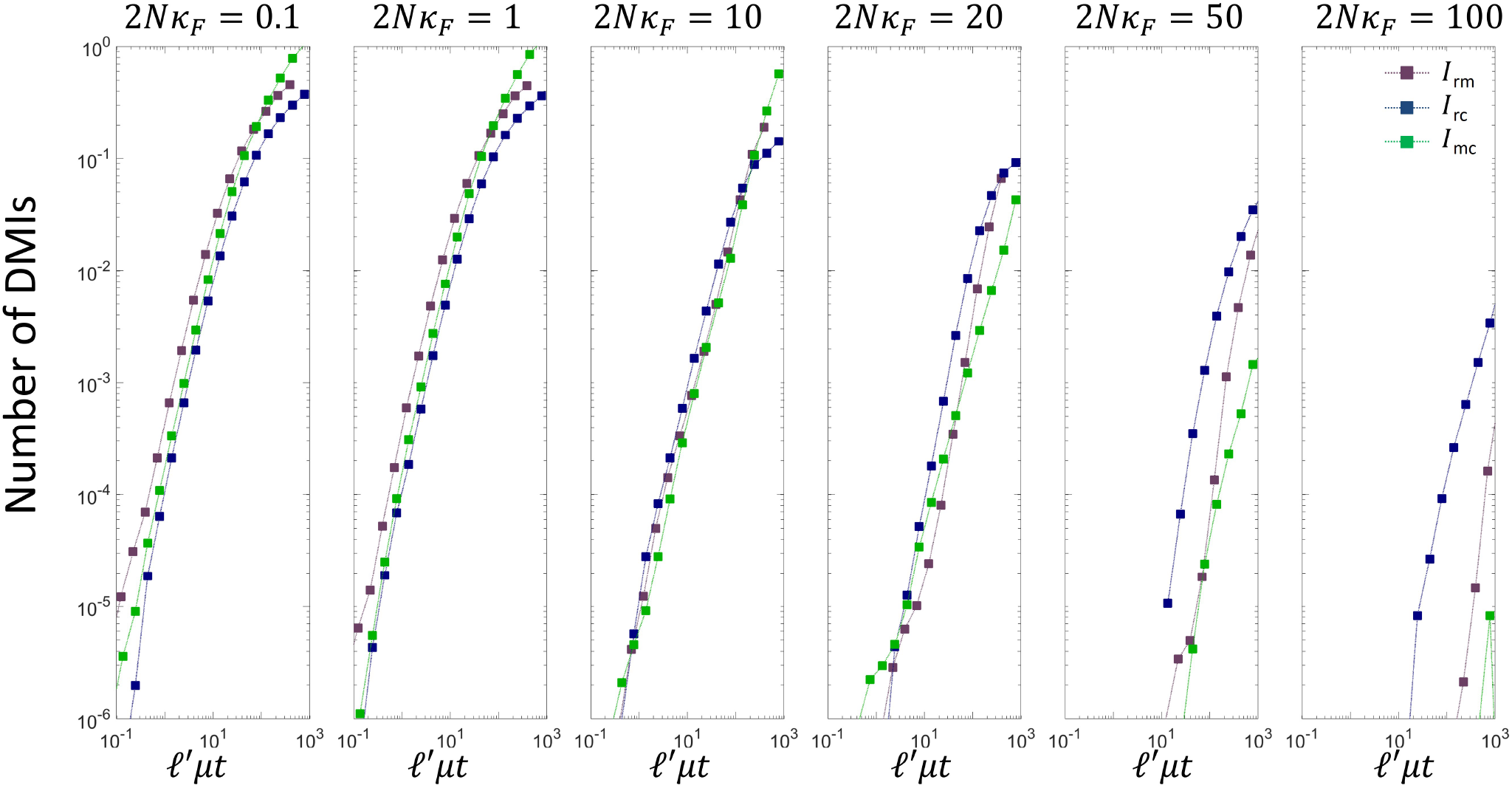
Plot of the spectrum 2-way DMIs vs scaled divergence time for different scaled population sizes. Here the divergence time for each pair-wise DMI is scaled by the number of base-pairs involved in each interaction in order to remove the effect that interactions with a larger number of base-pairs mutate at a larger rate in proportion to their length; for *I*_rm_, *ℓ*′ = 5, and for *I*_rc_ and *I*_mc_, *ℓ* = 10

In the converse limit, when population sizes are sufficiently small that all molecular phenotypes are neutral (up to the truncation threshold), we find only small differences between the rise of each of the pair-wise incompatibility types; this is again consistent with simple models of transcription factor DNA binding, where the phenotypic distribution of binding energies is dominated by their sequence entropy and not fitness at small population sizes and there is only a weak dependence on sequence length [43]. In the Supporting Information, we show this is also true of the higher order DMIs, where it is purely the sequence-entropy constraints of each DMI type which controls the dynamics of incompatibilities and fitness has little bearing; however, as the population size is increased this degeneracy is lifted and each of the DMIs exhibits individual behaviour.

## DISCUSSION

There is still very little understood about the underlying genetic basis that gives rise to reproductive isolation between lineages. Gene expression divergence is thought to be a strong determinant of the differences between species [1, 47, 82–84] with a growing body of evidence for their direct role in speciation, particularly through transcription factors [6, 10, 51, 53, 75]. In particular, transcription factors mediate the control of gene expression and the ultimately body plans of organisms through complicated gene regulatory networks. Studies in drosophila and nematodes have shown in closely diverged species conserved body plans yet plasticity in the underlying molecular architectures between them [26, 33, 76, 79].

This evidence suggests that divergence in the regulatory networks controlling gene expression and body plans in allopatric lineages is likely to be an important mechanism of speciation in many higher organisms, yet we do not yet have a quantitative evolutionary framework to model such processes, which can then be used to make predictions. A key challenge is to link changes at the genetic level, where mutations arise, to their outcomes on changes in phenotype, where selection actually acts. Although in the past century we have made great strides in our understanding of evolution through the modern synthesis of Fisher, Wright, Kimura and many others [15, 18, 20, 46, 85], this has focussed on the evolution of either genotypes or phenotypes separately. However, in recent years there has been increasing attention on the development of more realistic fitness landscapes, and how this affects the evolutionary process. For example, in biologically realistic systems, often many genotypes correspond to the same phenotype; this redundancy of the mapping from genotype to phenotype results in a number of non-trivial and emergent behaviours which do not arise on fitness landscapes which consider evolution of phenotypes or genotypes independently [7, 23, 29, 31, 32, 35, 44, 52, 55, 59, 69].

One important and well explored example is the evolution of transcription factor DNA binding [7, 34, 42, 43, 52, 59, 60], where the genotype-phenotype map from sequence to binding affinity can be explicitly enumerated under simplifying assumptions [27, 80]. These investigations show that for small populations dominated by genetic drift, evolution does not optimise fitness. Rather, there is a trade off between the high fitness of a small number of sequences that bind well and the exponentially larger number of sequences that bind less well. The result is the maximisation of a combination of fitness and the number of sequences that correspond to that phenotype. We can take advantage of analogies with statistical mechanics and represent this combination as the “free fitness”, where the log of the number of sequences is the “sequence entropy” of a phenotype [7, 37, 42, 43]. In this formulation, the effective population size is analogous to an inverse temperature for a physical system connected to a heat-bath, where decreasing population size increases the effect of drift and the importance of sequence entropy relative to fitness.

When the free fitness framework is applied to the role of transcription factor DNA binding in allopatric speciation, our previous work gave rise to a simple prediction: incompatibilities arise more quickly for smaller, drift-dominated, populations [42, 43], supporting previous computational studies by Tulchinsky et al. [77], that showed decreased hybrid fitness for smaller populations. This can be understood as a result of the greater importance of sequence entropy for small populations, resulting in common ancestors with a higher drift load, which are therefore closer to incompatible regions. As a result, fewer substitutions are required for the development of hybrid incompatibilities [42, 43]. Conversely, those transcription factor binding site pairs under weaker selection, at a fixed population size, will give rise to incompatibilities more quickly, as they are more susceptible to drift and in the common ancestor will have a larger drift load.

In this paper, we examine speciation in a more realistic genotype phenotype map. For the first time we examine how incompatibilities arise in allopatry for a simple evolutionary model of developmental system drift, where a higher level organismal spatial patterning phenotype is maintained by stabilising selection, whilst the underlying molecular binding energy phenotypes and ultimately the sequences that determine them, the genotype, are allowed to drift in the evolutionary simulations. Earlier analyses of this model demonstrated the evolution of a number of non-trivial features such as a balance between fitness and sequence entropy deciding the course of evolution at small population sizes and a roughness to the fitness landscape for phenotypes which have high fitness [44]. Importantly here, unlike in previous works [42, 43] we do not directly select for high binding affinity, but only on the organismal level phenotype, but as we discuss, we find the same population size dependence, as well as a number of other novel phenomenon to the speciation process, which would not be obtainable by modelling selection only at the level of phenotypes or genotypes. The results show that biophysics and effective population size provide a much stronger constraint than previous simple modelling of the dynamics of hybrid incompatibilities would suggest [64, 66].

A key result we find is that small populations are characterised by a power law growth of incompatibilities with time, vs large populations a sub-diffusive law (discussed below). Thus we suggest that empirical evidence of power law growth in incompatibilities is a signature of allopatric speciation at small population sizes. The Orr model of the growth of hybrid incompatibilities predicts that incompatibilities grow as a power law of the divergence time between allopatric lineages [64, 66], where the exponent represents the number of genes participating in the interaction (e.g. 2 for a 2-way incompatibility). The results of our model also yield this prediction, but only when populations sizes are sufficiently small. There is, however, an alternative model for the power law behaviour to the combinatoric argument made by Orr. As argued in [43], at small population sizes,where genetic drift is dominant and there is a large drift load, common ancestor populations are poised close to the incompatibility boundary and the growth of DMIs at short times is determined by the likelihood that a few critical substitutions arrive quickly, which is given by a Poisson process; if the critical number of substitutions is *K*\* then for short times we would expect *P_I_*(*t*) ~ (*μt*)^*K*\*^ and so given that at least *n* substitutions are needed for a *n*-way incompatibility, we would expect *K*\* ≥ *n*. In this paper, we introduced a new method to decompose DMIs into their fundamental pair-wise, 3- and 4-way incompatibilities, and find that for more complex incompatibilities (more loci involved) the larger the exponent of their power law growth. However, we find the exponents we measure for 3- and 4-way incompatibilities are smaller than the predicted exponents of 3 and 4 respectively. We suggest this could be due, as shown in the Supporting Information, to the greater number of higher order DMIs arising just by chance, leading to an overestimation of 3- and 4-way DMIs at short times, where at short times a *smaller* exponent corresponds to a larger number (i.e. *τ*^*n*−1^ > *τ^n^* for *τ* < 1, where τ is some dimensionless timescale).

We also find that incompatibilities arise more rapidly in smaller populations, which is an emergent effect due to the genotype-phenotype map, giving a bias in degeneracy of different phenotypes; lower fitness and less sharp patterning organismal phenotypes have many more sequences than higher fitness, sharper patterning, phenotypes. In smaller drift-dominated populations, this means there is bias towards phenotypes of small sequence entropy (log degeneracy) that counteracts the tendency of natural selection to favour phenotypes of high fitness. Consequently, the common ancestor in small populations is more likely to be slightly maladapted and less substitutions are needed before hybrids develop incompatibilities. These predictions are consistent with empirical evidence for an inverse correlation of speciation rates with effective population size; the net rates of diversification from phylogenetic trees [2, 14, 61] indicate smaller populations speciate more quickly, as well from in ferred times for post-zygotic isolation to arise [13, 21, 73], where for example mammals and cichlids, which have effective population sizes of order 10^4^ [22, 25, 40, 78], develop reproductive isolation more quickly than birds, which have effective populations sizes of order 10^6^ [68]. This model and the similar results obtained for transcription factor DNA binding [42, 43] provide a robust explanation of how stabilising selection can give rise to this population size effect in speciation, which do not require passing through fitness valleys as do models based on the founder effect [3, 4, 48, 49]. However, the results for this genotype-phenotype map for developmental system drift are particularly noteworthy compared to the previous results on transcription factor DNA binding, as they are obtained without directly selecting for high affinity, low sequence entropy, binding phenotypes; here we only select on the organismal spatial patterning phenotype, but nonetheless we find small populations develop hybrid incompatibilities more quickly through a similar mechanism of the interplay between fitness, sequence entropy and populations size. Although studies with more complex genotype-phenotype maps will be required, we suggest this points towards a broad principle, where the specificity required of a phenotype to be functional and of high fitness will mean that it will be coded by fewer genotypes. For example, in simple models of protein stability, the empirical observation that all proteins tend to be marginally stable, can be explained by the fact that as the stability of a protein is increased the number of sequences that give that stability decreases rapidly [29]; assuming high fitness corresponds to maximum stability, this phenotype is highly specified, as only a few sequences will meet the requirement that all inter-chain interactions in the protein are favourable.

Examining the growth of incompatibilities at large population sizes, we see there is a characteristic negative curvature on a log-log plot, predicted theoretically by [42], indicating that, as the number of substitutions needed for incompatibilities is large, the changes in the hybrid traits can be approximated by a diffusion process. However, we find that a simple model of diffusion does not fit the simulation data well; instead a model of sub-diffusion, that would arise if there are a number of kinetic traps giving a broad distribution of substitution times, does fit the data well. This is consistent with the finding that the genotype-phenotype map has a rough fitness landscape, which is only revealed at sufficiently large population sizes [44].

Another property that emerges from this model not obtainable by simply modelling transcription factor DNA binding is that certain molecular phenotypes are more important than others in giving rise to incompatibilities. One particular feature of this model is that the selective constraints on the different molecular binding energy phenotypes emerge through the evolutionary process of stabilising selection on the organismal phenotype, and are not specified by the model. Most strikingly, and counterintuitively, the model predicts that molecular phenotypes that are under the weakest selective constraints (but not strictly neutral) dominate by giving rise to the earliest incompatibilities for intermediate and large population sizes. Remarkably, here this emerges as a consequence of stabilising selection on the organismal phenotype and not due to selection imposed for good binding affinity as in previous works [43].

We note that these results have been obtained by changing the population size whilst keeping the strength of selection on the organismal trait *κ_F_* fixed. It would be tempting to use these results to suggest that overall those traits in a genome under weakest selection would give rise to the earliest incompatibilities and hence dominate allopatric speciation. However, the question of the how the dynamics of hybrid incompatibilities changes as the strength of selection changes is a subtle one, which we leave to future work; in this model a reduction in *κ_F_* has the effect of changing the phenotypic regions of incompatibility, with non-trivial consequences. It should also be noted, the role of sequence entropy in giving a strong population size dependence to the rate of reproductive isolation; if we consider only a peaked phenotypic landscape without a sequence entropy bias, a reduction in population size would only lead to a broadening of the phenotypic distribution and not a change in the mean of the distribution, and so a much weaker effect, as the common ancestor will still be *typically* far from incompatible regions. On the other hand with strong (exponential) degeneracy biases, the mean phenotype of the common ancestor changes strongly giving the large population size effect seen, which is as demonstrated in Fig.2.

Another finding of significance is that pair-wise or 2-way DMIs dominate compared with higher order DMIs (3- and 4-way in this model with 4 loci). This is in contrast to Orr’s theoretical argument that predicts a very specific ratio of 2-way: 3-way: 4-way DMIs, equal to 12 : 24 : 14, which assumes that the fraction of viable paths from the common ancestor to the current day species increases as we consider higher order DMIs [64]. This argument partly rests on the assumption that the number of inviable genotypes remains fixed as a larger number of loci are considered, which would seem a very strong assumption. Despite its simplicity, the genotype-phenotype map in this paper has many of the key features required for higher levels of epistasis, with protein-DNA binding, protein-protein binding and control of the morphogen steepness, all interacting in a nonlinear fashion to produce a single gene expression patterning phenotype and so there is clearly the potential for the Orr prediction to be verified; in contrast, we find the converse and our results show there is no bias towards 3-way DMIs, in fact showing instead that the ratio of 2-way to 3-way DMIs is at short times many orders of magnitude larger. This suggests, in this simple, but still relatively complex model, that biophysical constraints are far more important than a purely combinatorial argument would suggest. Evidence could be obtained from more detailed studies similar to [56, 57], where a power law with an exponent greater than 2 would indicate higher-order DMIs are dominant; currently this evidence suggests a quadratic growth law, however, a study with more time-points or species-pairs would provide more confidence. An alternative approach would be to look for linkage disequilibrium between unlinked regions of hybrid genomes, such as was found with hybrids of two species of swordtail fish [70], and though computationally challenging and requiring large numbers of parallel datasets, compare this against evidence for pervasiveness of higher order epistasis. Although recent results of [81], would seem to contradict our conclusions, their finding of extensive complex epistasis relates to higher order interactions between sites within a single locus, coding for protein stability or enzymatic activity, whereas our work relates to epistasis between multiple loci. Similarly, the results of hybrid incompatibilities within RNA molecules [41], which show a ‘spiralling complexity’ of DMIs would appear to be of limited biological relevance to allopatric speciation in higher organisms, as these are related to epistasis within a single locus, which are unlikely to segregate into different recombinants in a hybrid.

Finally, for small populations we find clustering in the behaviours of growth of different types of DMIs, in particular, 3-way DMIs (Supporting Information), which can be explained by the different sequence entropy constraints on different molecular phenotypes. This degeneracy is then lifted at larger population sizes and each *n*-way DMI takes on a different identity in their pattern of growth; this has strong analogy to physical phenomenon in statistical physics where constraints of symmetry dominate at large temperatures, in a regime where noise is important, but at smaller temperatures this symmetry is broken.

A clear future direction to investigate would be the effect of multiple transcription factors binding to enhancer regions to control gene expression [9, 50, 72] in large gene networks, where there is potential scope for complex epistasis across many loci coding for a large number of transcription factors. However, as our results show, despite the possibility and a prior expectation of a larger number of triplet interactions, pair-wise interactions dominate; for complex transcriptional control, if pair-wise interactions between proteins, and proteins and DNA dominate, for example in determining the binding affinity of transcriptional complexes, then our conclusions would hold. As we broaden the scope to large gene regulatory networks, there is no strong and direct empirical evidence for pervasive higher order epistasis in their function, which could give rise to higher order incompatibilities being dominant [74]. Specifically, although there is evidence that higher order incompatibilities have arisen in natural populations [12, 24, 65, 67], nonetheless a survey of these findings suggest there is no evidence for their dominance [24] as would be predicted by Orr and would be consistent with our findings that point towards biophysics providing a stronger constraint.

Overall, our results point to a basic principle, where developmental system drift or cryptic variation [26, 33, 76, 79], play a key role in speciation; basic body plans or phenotypes are conserved, but co-evolution of the components and loci of complicated gene regulatory networks can change differently in different lineages, giving incompatibilities that grow in allopatry. Here, we suggest a universal mechanism, where the rate of growth of incompatibilities is controlled by the drift load, or distribution of phenotypic values, of the common ancestor, which in turn is determined by a balance between selection pushing populations towards phenotypes of higher fitness and genetic drift pushing them towards phenotypes that are more numerous (higher sequence entropy); this basic mechanism would predict in general that traits under weaker selection will dominant the initial development of reproductive isolation. In particular, although in principle more complicated regulation could give rise to more complex patterns of epistasis [64], our findings suggest that more simple, pair-wise, incompatibilities dominate the development of reproductive isolation between allopatric lineages under stabilising selection.

## METHODS

### Details of the GP map

We model the binding energies of proteins to DNA using the “two-state” approximation [27, 80], which assumes that the binding energy of each amino acid-nucleotide interaction at the binding interface is additive and to a good approximation controlled by the number of mismatches, which each have the same penalty in binding affinity. The various protein-DNA binding energies in the main text are then given by the Hamming distance between the respective sequences. For example, the binding energy between the morphogen (*M*) and the first binding site (*B*) is given by

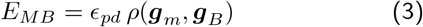

where *ρ*(***g**_m_*, ***g**_B_*) is the Hamming distance between the protein binding sequence (***g**_m_*) for the morphogen and the sequence for a first regulatory binding site (***g**_B_*), where *ϵ_pd_* is the cost in energy for each mismatch. We assume *ϵ_pd_* = 2*k_B_T* as a typical value for the mis-match energy, which are found to be in the range 1 − 3*k_B_T* [27, 80]. Similarly the co-operative protein-protein binding energy, for example between RNAP and the morphogen is *Ẽ_RM_* is

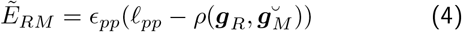

where ***g**_R_* is the sequence involved in protein-protein interactions for the polymerase, and 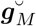 represents the equivalent binary sequence for the morphogen, flipped about its centre, which mimics the chirality of real proteins and prevents the co-operative stability from always being maximum for homo-dimers. The parameter *ϵ_pp_* is the stability added for each favourable hydrophobic interaction between amino acids, which we assume to be *ϵ_pp_* = −*k_B_T*. Given *ℓ_pp_* = 5 this gives interactions consistent with typical literature values of −2 to −7*k_B_T* for hydrophobic interactions between proteins [11, 71].

The morphogen concentration profile is approximately exponential; the exact profile we use is

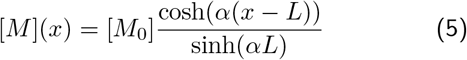

where this arises from solving the reaction-diffusion equation with reflecting boundary conditions and is valid for all *α*.

### Monte Carlo scheme for speciation simulations

We use a kinetic Monte Carlo scheme to simulate the evolutionary process for the genome ***G*** and *α* on two independent lineages, assuming a fixed effective population size of *N*, and that we are in the regime of small effective population size (*ℓ_**G**_μ*_0_*N* ≪ 1, where *μ*_0_ is the base-pair mutation rate). This means the population is represented by a single fixed sequence (or number for *α*) for all of the loci at each time-point, where effectively mutations are either instantly fixed or eliminated. Specifically, we use the Gillespie algorithm [28], to simulate evolution as a continuous time Markov process; at each step of the simulation the rate of fixation of all one-step mutations from the currently fixed alleles (wild type) is calculated, and one of these mutations is selected randomly in proportion to their relative rate. Time is then progressed by *K*^−1^ ln(*u*), where *K* is the sum of the rates of all one-step mutants and *u* is a random number drawn independently between 0 and 1, which ensures the times at which substitutions occur is Poisson distributed, as we would be expected for a random substitution process. The rates are based upon the Kimura probability of fixation [45]:

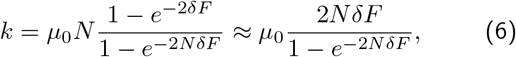

where *δF* is the change of fitness of a mutation at a particular location, given by fitness function defined in the main text, and *μ*_0_*N* is the rate at which mutations arise in the population; the latter approximation in Eqn.6 assumes *δF* ≪ 1. Note that although in the simulations we use the full form for the fixation probability, fitness effects are typically small (*δF* ≪ 1) in the simulations, so the substitution rates only depend on the population-scaled fitness changes 2*NδF* which, for a given mutation, is proportional to 2*Nκ_F_*. Finally, we allow continuous ‘mutations’ in the morphogen steepness parameter *α*, chosen from a Gaussian distribution of standard deviation *δα* = 0.5 and assign it an 10 effective base-pairs, which are used when assigning relative weight in the kinetic Monte Carlo scheme, where the total number of base-pairs is *ℓ_**G**_* = 60.

We determine the Malthusian or log fitness of the spatial gene regulation, from the resulting concentration profile [*TF*](*x*) by use of a functional that promotes expression of the TF in the anterior half, whilst penalising expression in the posterior half, with truncation selection below a critical value *F*\*:

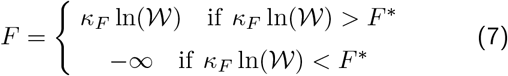

where,

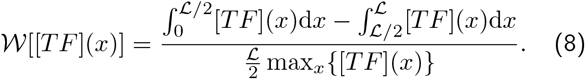

where we use a value of *F*\* = −1.6 × 10^−3^, which corresponds a value of 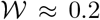, when *κ_F_* = 10^−3^. Strictly, an inviability on a lineage or a hybrid should correspond to 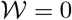 or *F*\* = −∞, however, these values were chosen to so that a reasonable number of incompatibilities arise in a simulation; for comparison the typical maximum of the integral 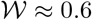. In this paper, we explore how the changing the population scaled strength of selection (2*Nκ_F_*) affects the rate of reproductive isolation, by keeping *κ_F_* fixed and varying *N* accordingly. Note that although here the exact form of the fitness is slightly different to the one used in [44], the qualitative behaviour is the same (Supporting Information).

The speciation simulations consist of two replicate simulations starting with the same common ancestor and with the same fitness function. We draw the common ancestor from the equilibrium distribution for ***G*** and *α*. To do this we start from a random initial genome, and run one long simulation for 100,000 substitutions for a fixed scaled population size 2*Nκ_F_*, in order to effectively equilibrate the system (typically 10,000 substitutions are required to adapt to an ensemble of fit states). This represents a reference equilibrium state; different random draws from the equilibrium distribution then consist of running the simulation for a further 100 substitutions.

### Decomposing DMIs

Given a pattern of hybrid incompatibilities, for example, as shown in Fig. 4, if there is a 2-way DMI (e.g. between *C* and *α* loci, which we denote *I*_Ca_), then all four hybrid-genotypes containing this DMI (e.g. RMCa, RmCa, rMCa, rmCa) are inviable; these genotypes define a twodimensional subspace (or face) of the hypercube. Similarly, the points (e.g. rmcA, RmcA, which we denote *I*_mcA_) containing a 3-way DMI form a one-dimensional subspace (or line), while a 4-way DMI takes up only a single point in the four-dimensional hypercube. These different 2-way, 3-way and 4-way DMIs are the fundamental incompatibility types which we seek to explain the pattern of hybrid inviable genotypes observed, for example, as in Fig.4a.

However, this decomposition is hugely underdetermined, as there are only 2^4^ − 2 = 14 possible hybrids (not including the well-adapted genotypes of line 1 and line 2) and a total of *I_max_* = 3^*L*^ + 1 − 2^*L*+1^ = 50, different fundamental incompatibilities, for *L* = 4 loci. This arises as the total number of *n*–point DMIs is 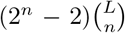, as there are 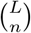 combinations of *n* loci amongst *L* total loci and then considering a binary choice of alleles across both lines, there are a total of 2^*n*^ allelic combinations or states, 2 of which are the fit allelic combinations where all alleles come from one lineage or the other giving 2^*n*^ − 2. For example, between each pair of loci there are 2^2^ − 2 = 2 mismatching combinations of alleles (e.g. rMand Rm) that could give DMIs and 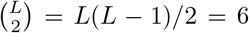 pairwise interactions. A similar argument would give a total of 24 3-way DMIs as there are 2^3^ − 2 = 6 mismatching combinations of alleles at 3 loci (e.g., excluding rmcand RMC) and 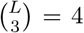 3-way interactions and similarly, 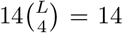 for 4-way interactions. In total, the max number of DMIs is 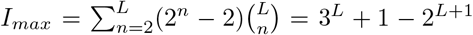 which for *L* = 4 loci is *I_max_* = 50.

The approach we take is to find only those combinations of fundamental DMIs that have the smallest total number that can explain the pattern of hybrid incompatibilities, which from a Bayesian perspective would have the smallest Occam factors [54]; for instance, as shown in Fig.4b the list of 6 incompatible hybrid genotypes rmCa, rMCa,RmCa,RMCa,Rmca,RMca, shown by red crosses, can be explained most parsimoniously by three different minimal combinations of fundamental DMIs, each with only 2 DMIs (see main text).

## ACKNOWLEDGMENTS

We would like to thank Davis Pollock for initial discussions. RAG was supported by the Medical Research Council under grant U117573805 and BSK by The Francis Crick Institute which receives its core funding from Cancer Research UK, the UK Medical Research Council and the Wellcome Trust.

## Supporting Information

### EMERGENT PROPERTIES OF GENOTYPE-PHENOTYPE MAP

The properties of this genotype-phenotype map have been previously explored [S44]. So here we summarise its pertinent findings. Under a fixed environment (fixed fitness) adaptation occurs in a manner which is insensitive to initial conditions and to the same ensemble of states[S62]. As shown in Fig.S1, the evolutionary simulations find solutions to the patterning problem, where it is clear that for a small scaled population size (2*Nκ_F_* = 1) the solutions fixed at any one time are less constrained, while as the population size increases (2*Nκ_F_* = 50) the solutions are more constrained. This is reflected in the time series of the fitness as shown in Fig.S2a, where we see that for 2*Nκ_F_* = 1 the fitness of solutions are low, with large fluctuations and very often near the inviability threshold *F*\*, whereas for 2*Nκ_F_* = 50 the fitness is higher and far more constrained. We see this also in the time series of the phenotype *α* (Fig.S2b), which shows that it is small with large fluctuations for 2*Nκ_F_* = 1 and is larger and more constrained for 2*Nκ_F_* = 50, which reflects the fact that a larger value of *α* corresponds to a more steep gradient of the morphogen, which in turn allows sharper patterning.

As was found in [S44], there are a large number of genotypic states that give good patterning and they all belong to the same solution to the patterning problem; the morphogen binds cooperatively with the polymerase to promote transcription when the morphogen concentration is large and turn off transcription when the morphogen concentration is small. As demonstrated by the histograms of the binding energy phenotypes in Fig.S2c&d, this solution in general requires *E_MB_* and *Ẽ_RM_* to be small (strong binding) and that *E_RP_* isn’t too small, so that a switch-like mechanism exists to allow the polymerase to be attracted to the promotor in the presence of a large concentration of morphogen. The exact state of G determines these binding energies and then *α* is determined by the requirement that the threshold morphogen concentration which switches on or off transcription of *TF* occurs at the mid-point of the embryo. It is clear from Fig.S2c&d that some of the energy phenotypes are less important than others, in particular, *E_MP_, E_RB_, Ẽ_MM_* & *Ẽ_RR_* all have broad peaked distributions, even for large populations sizes, consistent with the neutral distribution of the binding energies, given this Hamming model, following a binomial distribution peaked about approximately about *ℓϵ*/2, and so indicating that the fitness constraints on these phenotypes is relatively weak. Finally, an emergent property of the solutions is that there is a division of genotype space into a high sequence entropy region, where there are many sequences that code for slightly sub-optimal solutions corresponding to a small value of *α*, and a low sequence entropy, higher fitness region, which corresponds to a large value of *α* − this is what gives rise to the bistability visible in *α* for 2*Nκ_F_* = 50 (see [S44] for more details). On long timescales, we see transitions between these regions, whilst on short timescales we mainly see exploration of each of these basins of attraction.

### DIFFERENT HYBRID GENOTYPES HAVE DIFFERENT GROWTH RATES OF DMIS

To examine in more detail the overall trends in the number of incompatibilities for each hybrid type, we plot the average number of DMIs as a function of divergence time *μt* in Fig.S3, where we have averaged over pairs of complementary genotypes (e.g. RmcA and rMCa), which have identical statistical properties. In Fig.S3, we first note that each hybrid-genotype behaves in a different, population size dependent manner. In general, there is an initial growth in the average number of DMIs as the divergence time increases, followed by a plateau or a slowing down of the growth. As denoted in the figure, we distinguish hybrid-genotypes based on the types of potential mismatch: *R*-type is characterised by a mismatch of the *R* locus with the *M* and *C* loci, *M*-type is a mismatch of *M* with the *R* and *C* loci (e.g., RmCa), *C*-type is a mismatch of *C* with *R* and *M* (e.g. rmCa) and *α*-type is a mismatch of the *α* locus with the rest of the loci. For each of the *R*-, *M*- and *C*-types, we assume the interaction with the *α* locus is relatively weak, so or example, the hybrid genotype Rmca or RmcA are both *R*-type.

**FIG. S1.**
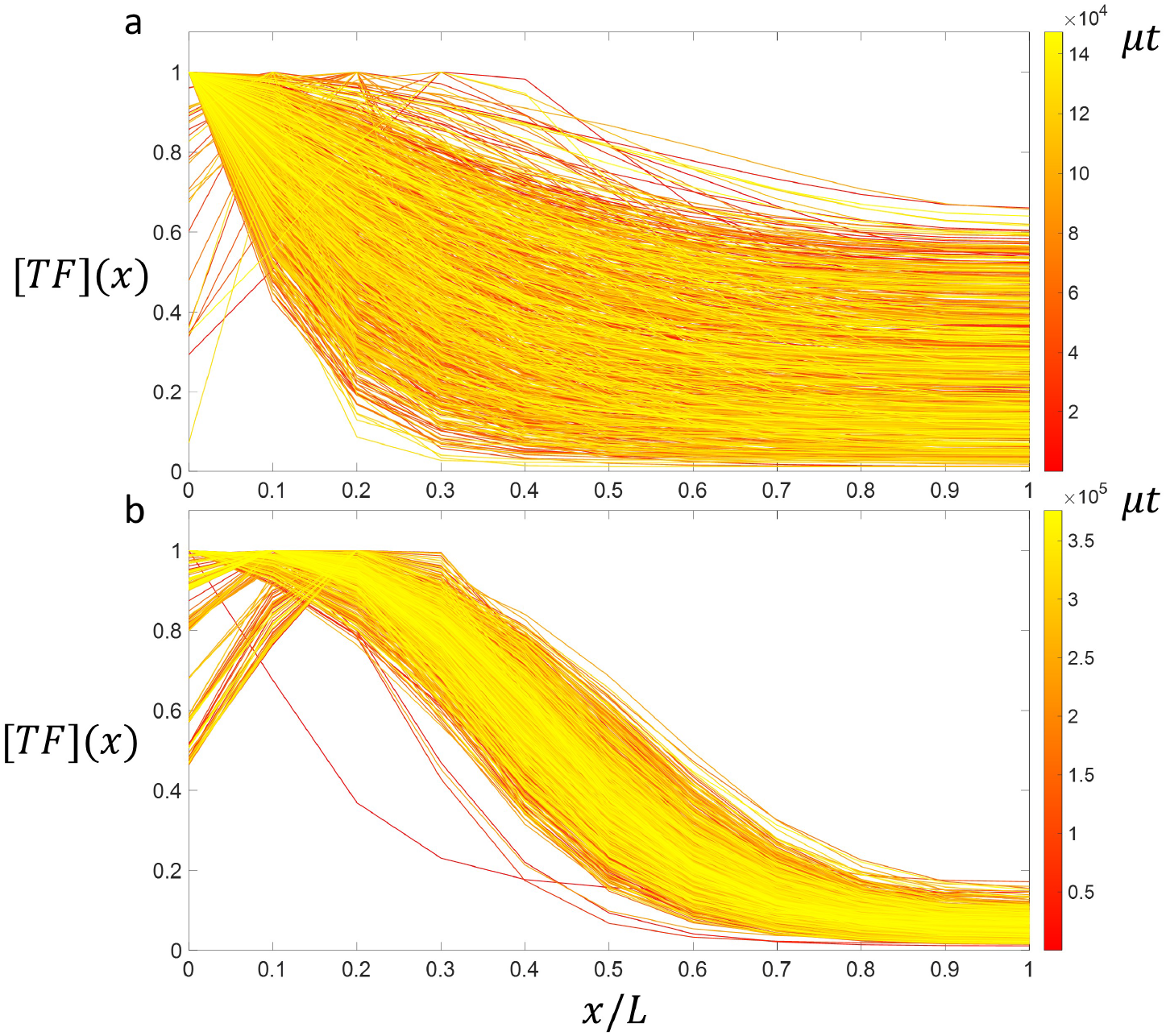
Typical spatial patterning profile of the output transcription factor [*TF*](*x*) over the course of the evolutionary simulations, where early times are indicated by red, becoming progressively yellow at late times (as indicated by the colour bars); a) large population size 2*Nκ_F_* = 50 and b) smaller population size 2*Nκ_F_* = 1. the results show evolution gives solutions which are less well adapted.

In addition to the general trends pointed out in the main text, we see that the growth of DMIs is different for different hybrids; for small population sizes (2*Nκ_F_* = 0.1 & 2*Nκ_F_* = 1), *M*-type, *C*-type and *R*-type dominate the growth of DMIs, in this order and with only a small difference between them, whilst *α*-type arise far more slowly; we might expect this since substitutions in *α* only tend to shift the pattern away from the mid-point of the embryo, which with the model of fitness defined in Eqn.7 only moderately affects fitness. As the population size increases, and at small times, we see that initially *M*-type DMIs arise more slowly relative to *C*-types and *R*-types, but at longer times there is a cross-over where *M*-type DMIs dominate *R*-type, which moves to longer times at increasing population sizes.

How can we understand this general behaviour? It is clear from Eqns. 7&8 that we need co-evolution of the relevant sequences to maintain these the important binding energies within certain constraints; e.g. the binding energy of *R* to the promoter *P, E_RP_* mustn’t be too strong and so on each line the sequences will co-evolve to maintain this constraint. From previous work on transcription factor DNA binding [S42, S43] we expect that the rate at which incompatibilities arise due a particular transcription factor binding site interaction *j*, is controlled by the product of the population size and the relative selection coefficient *κ_j_* (which should be distinguished from the overall selection coefficient for the patterning phenotype *κ_F_*); so the binding energy *E_MB_* is under very strong selection to give high binding affinity of the morphogen to the 1st binding site, whilst for example, the binding energy *E_RP_* is under weaker selection and so as *κ_RP_* > *κ_MB_*, we would expect incompatibilities due to this interaction to arise more slowly when 2*Nκ_MB_* ≫ 1. Similarly, since *κ_MB_* > *κ_j_*, for all other interactions *j*, we would expect *M*-type hybrids to develop incompatibilities more slowly at scaled large population sizes, as observed in Fig.S3.

**FIG. S2.**
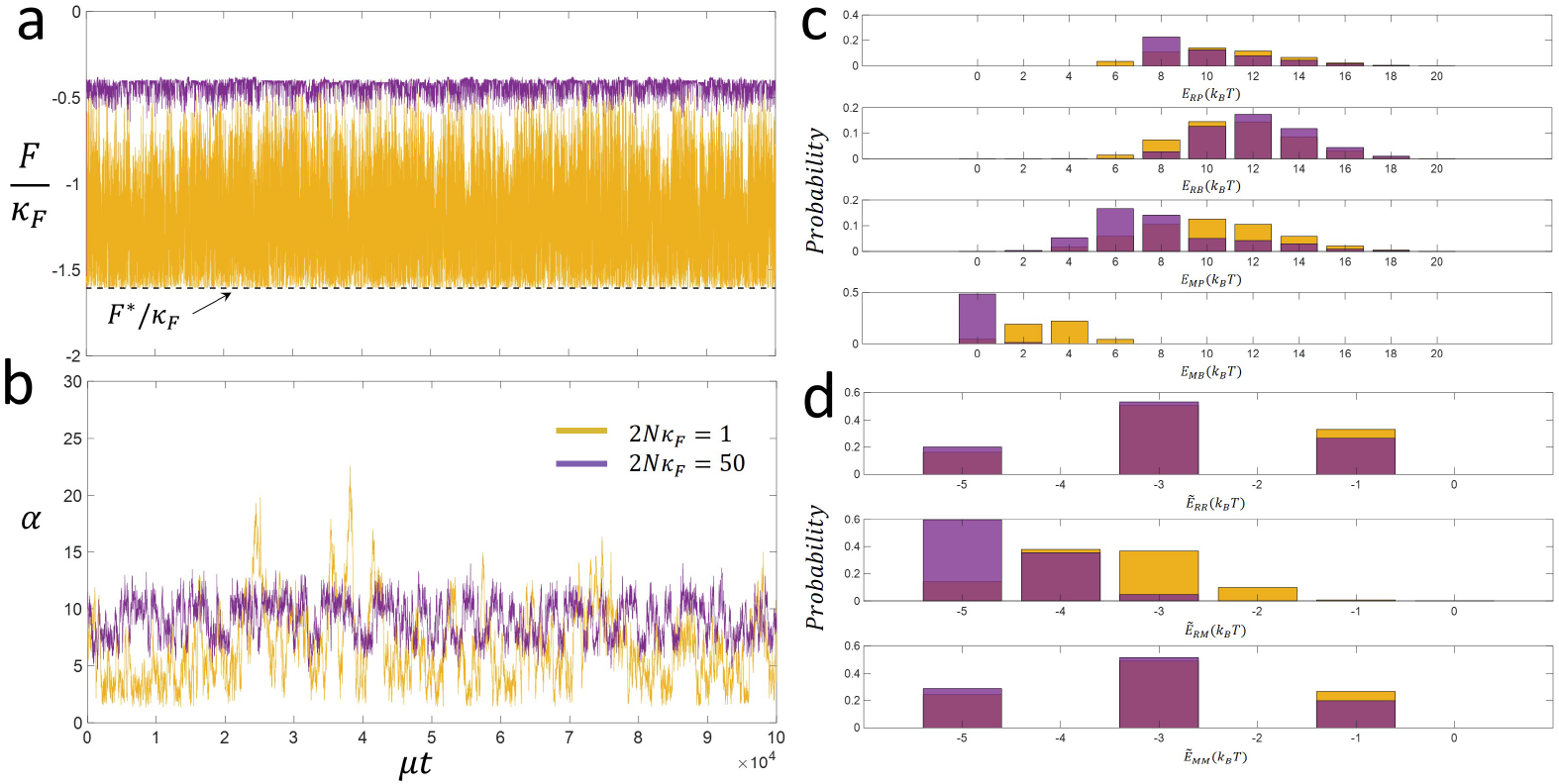
FIG. S2. Equilibrium properties of phenotypes and fitness of genotype-phenotype map at the population sizes of 2*Nκ_F_* = 50 (purple) and 2*Nκ_F_* = 1 (yellow); a) Malthusian fitness (defined by Eqn.1&2 in the main text) over time, where we see whereas at large population sizes fitness is near the optimum, at low population sizes the fitness of solutions are sub-optimal and very commonly near the inviability boundary for functional vs non-functional spatial patterning; b) the morphogen steepness trait *α* where we see there is generally a broad variation at small population sizes, against more constrained variation at the larger population size; histogram of c) protein-DNA binding energy traits and d) of protein-protein binding energy traits, where we see that different binding energies are constrained to greater or lesser extent (indicating their selective or functional importance), which are further relaxed at smaller population sizes, indicating the increasing dominance of drift. In particular, as binding energies are determined by a sum of mismatches, the neutral distribution is a binomial distribution centred on *ϵ_pd_ℓ_pd_*/2 or *ϵ_pp_* + 1)/2, so that deviations from this distribution are a measure of the selective constraint.

### NULL SPECTRUM OF DMIS

To test whether the parsimony decomposition of DMIs observed in Fig.5 of the main text, where 2-way DMIs are dominant, is not due to an inherent bias to detect 2-way DMIs of the method, we applied it to the case of randomly assigning incompatibilities to the different hybrid genotypes. This is easily accomplished by drawing random binary numbers from the Bernoulli distribution with a probability p that any hybrid is incompatible. For each random collection of incompatible hybrid genotypes the parsimony method is applied and the number of each type of *n*-way DMI is counted (as detailed in the Material and Methods of the main text); this is repeated 1000 times to calculate the average number of *n*-way DMIs shown below. To allow some comparison to the results of Fig. 5 of the main text, the value of *p* is varied according to the average fraction of incompatible hybrids found at different time-points from the simulations in the main text. Hence, the panels of Fig.S4 correspond to the number of n-way DMIs expected at different time-points for different scaled population sizes, where p is mapped into time using the sum of all curves in Fig.4 of the main text to calculate the fraction of incompatible hybrids at each time point. This null spectrum shows the opposite behaviour to that found in the simulations in the main text demonstrating that the results observed are not due to a random bias to find 2-way DMIs using the parsimony method.

**FIG. S3.**
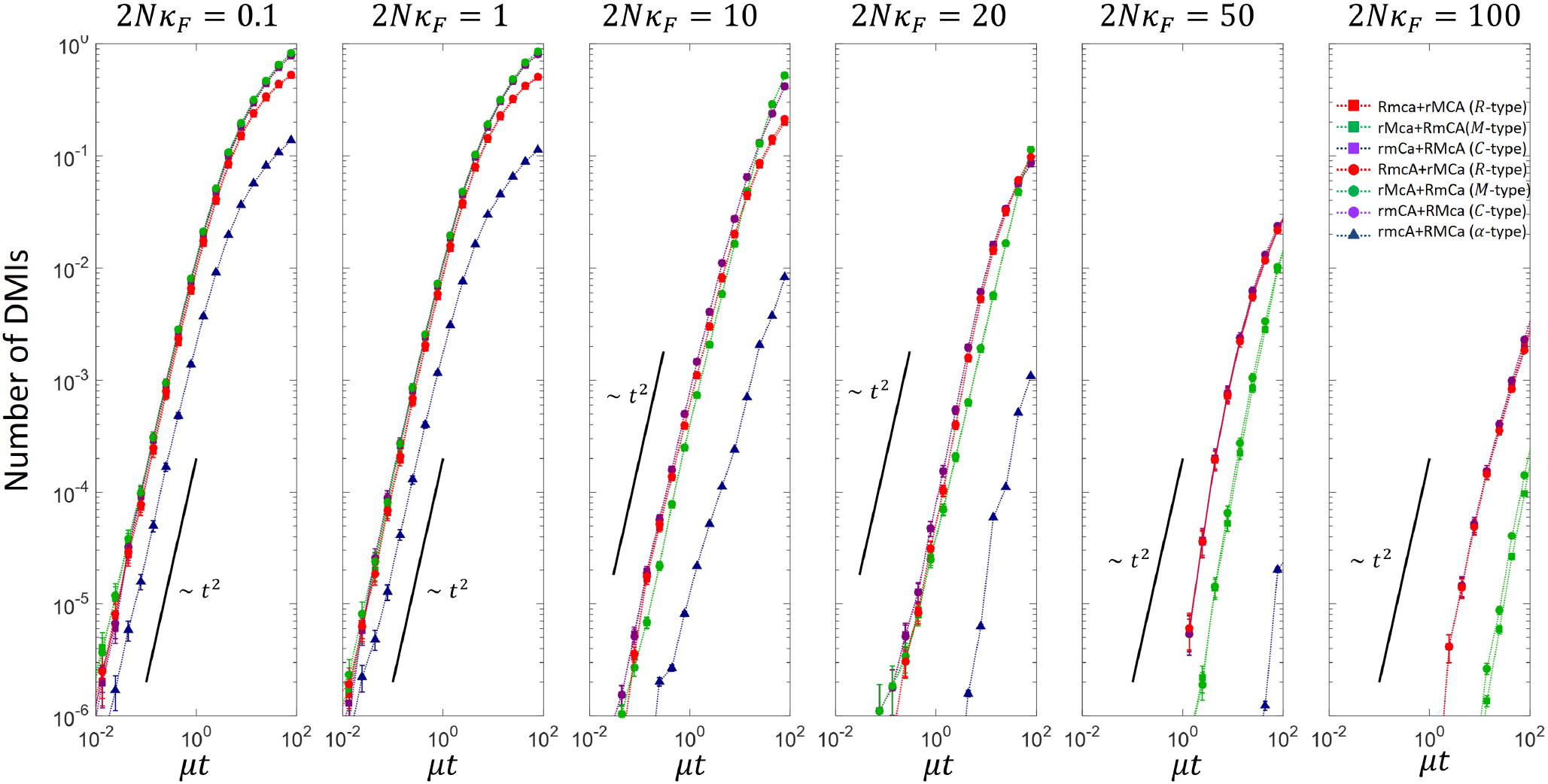
Plot of the number of DMIs for each hybrid genotype since divergence for different scaled population sizes. Complementary hybrid genotypes are summed over, since they have identical statistical properties.

### SPECTRUM OF DMI TYPES

#### Spectrum of 2-way DMIs

In Fig.S5 we have plotted the number of 2-way DMIs of each type, where for example, *I*_mc_ is a 2-way DMI caused by an incompatibility between the *M* locus and *C* locus. First, we note that at small population sizes the rate of increase of the different 2-way incompatibilities seem to cluster into two types; those that involve *α* and those that do not, where the latter arise more rapidly; this suggests that sequence entropy constraints are dominating for small populations, particularly at short times, though for the latter group there are some differences as discussed below. As the population size increases, we see that differences arise in the rate of growth between these different types of 2-way incompatibilities. Below we discuss these properties.

For each type of 2-way incompatibility there are 2 binding energy traits that could contribute. So increases in *I*_mc_ could be due to an incompatibility in the hybrid that can be traced to *E_MB_* or *E_MP_*; in this case, as the binding energy *E_MP_* is almost neutral [S44], we would expect incompatibilities to arise predominantly from *E_MB_*. Similarly, we expect *I*_rc_ to be dominated by *E_RP_* and not *E_RB_* and *I*_rm_ dominated by *Ẽ_RM_* and not *Ẽ_MM_* or *Ẽ_RR_*, as *E_RP_* is under strongest selection of the traits coded by the *R* & *M* loci. When it comes to incompatibilities involving the *α*-locus, there is no clear phenotypic trait that can be iden tified and as we will see the analysis of these DMIs will not be so clear.

**FIG. S4.**
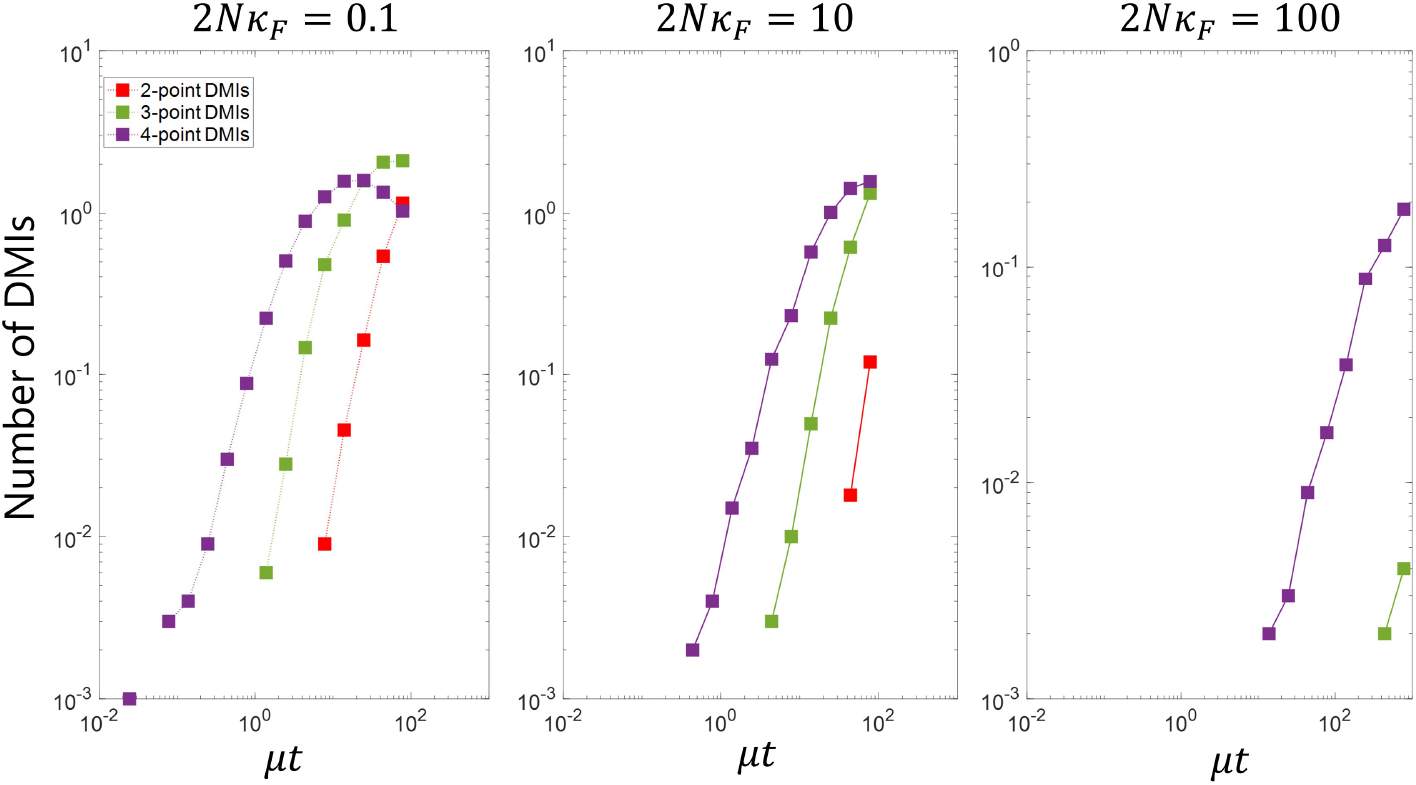
Parsimony decomposition of DMIs when incompatibilities are assigned randomly with a probability *p* that changes with time according to the fraction of hybrid genotypes that are incompatible from the simulations in main text.

**FIG. S5.**
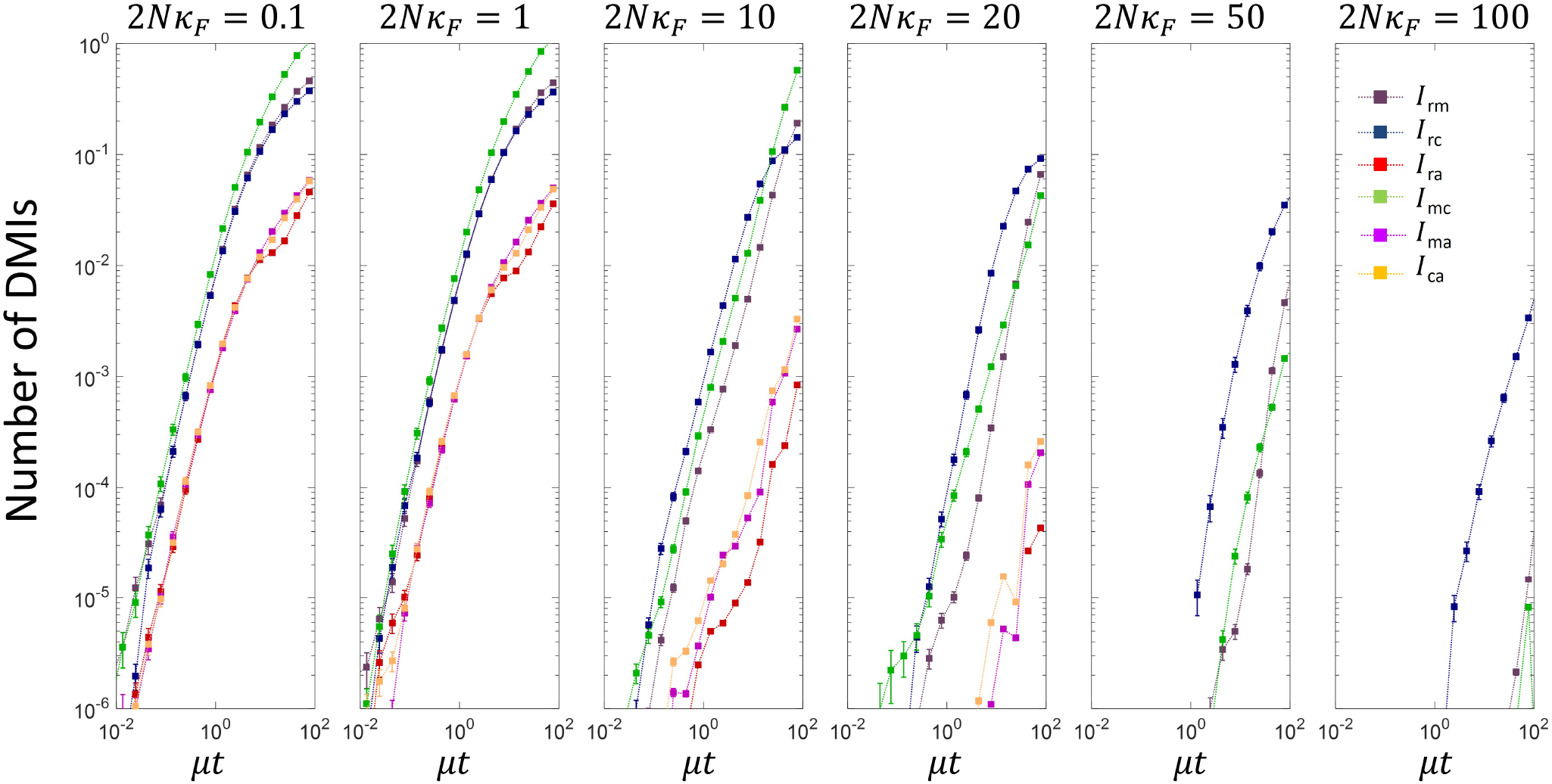
Plot of the spectrum 2-way DMIs vs divergence time for different scaled population sizes.

Examining Fig.S5, we see that as the scaled population size increases, the time for *I*_mc_ incompatibilities to arise sharply increases, while the time for *I*_rm_ increases less rapidly and *I*_rc_ even less rapidly. This corroborates a key prediction of a simple model of transcription factor DNA binding described in [S43], where traits under greatest selection should generate incompatibilities less rapidly, since the distribution of their trait values are further away from inviability in the common ancestor. As observed above, *E_MB_*, which contributes most to *I*_mc_, is under the greatest selection pressure and so as the population size changes these should change most rapidly. So we see that for large population sizes, it is not the phenotypic traits under the strongest selection that give rise to significant DMIs at short times, but those under a weaker selective constraint; traits under weaker selection will be affected more by the sequence entropic pressure for poorer binding affinities and so the common ancestor is more likely to be closer to the inviability boundary [S63].

However, at small population sizes, 2*Nκ_F_* ≤ 1, all DMIs grow approximately quadratically at short times with a saturating form at long times, as also seen in Fig.5. For small population sizes, the simple model of TF-DNA binding [S43] predicts that for the same sequence length and assuming the same threshold of inviability that the rate that incompatibilities arise should be approximately independent of selection and dominated by sequence entropy; if the sequence length of the trait increases, or the effective region of incompatibility is smaller, then we would expect incompatibilities to arise more quickly. As we see in Fig.S5, for small scaled population sizes and short times, the incompatibilities not involving *α* have similar behaviour except *I*_mc_, which arrive more quickly; this is as expected as the effective mutation rate for *I*_mc_ to arise, will be double that for *I*_rm_, since the binding of the morphogen to DNA involves 10 sites (nucleotides), vs 5 sites for protein-protein interaction. On the other hand *I*_rc_ also has 10 nucleotides, but arises more slowly. We suggest this could be due to a different effective region of incompatibility, which could confound simple expectations, it is not clear how a single inviability threshold *F*\* effectively maps to these pair-wise incompatibilities, complicating the picture further.

The 2-way incompatibilities involving the *α* locus are more difficult to interpret, since there is no clear trait in the patterning model associated with an interaction solely between the *α* locus and *R, M*, or *C* loci. As 2-way DMIs involving *α* are typically an order of magnitude smaller than the other DMIs they do not have a large impact on the number of DMIs.

#### Spectrum of 3-way & 4-way DMIs

In Fig.S6, we have plotted the 3-way DMIs as a function of divergence time *μt*, where the panels from left to right represent increasing scaled population size. We see that for small population sizes, 3-way DMIs between the *R, M* and *C* loci dominate at all times and in particular that the different types of DMIs of this type are all roughly equal, *I*_Rmc_ ≈ *I*_rMc_ ≈ *I*_rmc_. In addition, we see that all other DMIs arise more slowly and each of the 9 other types of 3-way DMIs are all again approximately equal. However, at larger population sizes this degeneracy is lifted amongst the different types of DMIs and different 3-way DMIs grow at different rates. How can we understand this general behaviour?

The patterning solution found in these evolutionary simulations involves the morphogen binding strongly to the first binding site recruiting the polymerase to bind to the promotor to turn on transcription, through a high affinity interaction between the morphogen and the polymerase; the spatial position along the length of the embryo where the transcription switches from on to off is controlled by an interaction with the steepness of the morphogen gradient *α*. Given this, incompatibilities between *R, M* and *C* loci could arise through a 3-way interaction where the *R* locus interacts with the parts of the *C* and *M* loci coding for *E_RP_* and *Ẽ_RM_*, or where the *M* locus interacts with the parts of the *C* and *R* loci coding for *E_MB_* and *Ẽ_RM_*. So in analogy to 2-way DMIs, where a pair of loci give rise a single phenotypic binding energy trait, whose value contributes to fitness, here the triplet of loci, *R, M* and *C*, give rise to two binding energy traits, which together contribute to fitness. These two traits will co-evolve to maintain good fitness, balanced by the constraints of sequence entropy on the underlying 3 loci; at large population sizes, the effects of sequence entropy will diminish. On the other hand 3-way incompatibilities between, for example, *M, C* and *α* could arise due to an interaction of the *E_MB_* binding energy trait with *α*; in this model this is subject to a sequence entropy constraint between only two loci. This is true for all the 3-way interactions that involve the *α* locus. Qualitatively, this then explains the behaviour at low population sizes, as sequence entropy dominates fitness, meaning that the behaviour of the different 3-way DMIs will be dominated by their underlying sequence entropy constraints.

**FIG. S6.**
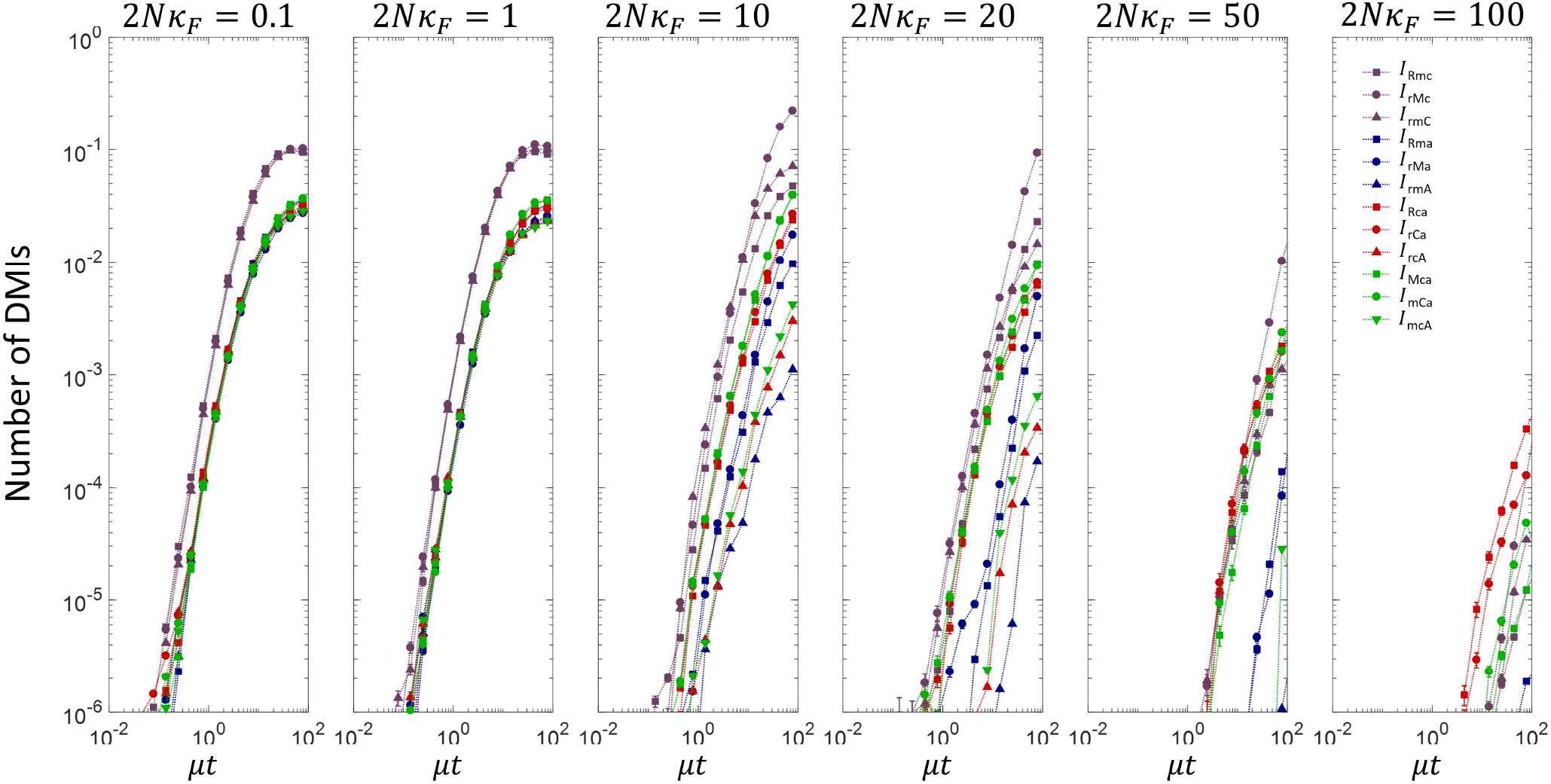
Plot of the spectrum 3-way DMIs vs divergence time for different scaled population sizes.

The sequence entropy constraints for the 3-way interactions involving the α locus is straightforward and given by a binomial degeneracy function 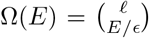, so that the sequence entropy function *S*(*E*) = ln(Ω) is approximately quadratic in *E*, where here *E* represents one of the binding energies that interacts with *α* [S42]. However, for the other 3-way interactions that don’t involve *α*, but involve the *R, M* and *C* loci, the sequence entropy constraint will be related to a degeneracy function Ω(*E, Ẽ*) = Ω(*E*)Ω(*Ẽ*) where the joint number of sequences that give E and E is a product, since these energy traits are coded by different sequences, even though they come from the same loci (the joint number of sequences Ω(*E_MB_, E_MP_*) ≠ Ω(*E_MB_*)Ω(*E_MP_*) since the protein binding sequence of the morphogen that determines *E_MB_* and *E_MP_* is the same in this case). Given that the joint number of sequences that give *E* and *Ẽ* is a product of two binomial coefficients, the sequence entropy function will approximately be a sum of two quadratic terms 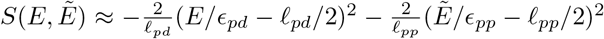. At small population sizes, where genetic drift dominates selection, we expect the distribution of common ancestors to be such that they are poised at the incompatibility boundary for *E* and *Ẽ*; incompatibilities then arise when substitutions arise that take hybrids across the boundary.

Given that a 3-way DMI between the *R, M* and *C* genetic loci corresponds to co-evolution of a pair of binding energy traits, instead of a single binding energy trait for 2-way and 3-way DMIs that involve *α*, means the fraction of substitutions that lead to incompatibilities versus those that keep the hybrids compatible/fit becomes larger when going from one to two dimensions. This then explains why 3-way DMIs between the *R, M* and *C* loci gives rise to incompatibilities more quickly than those involving the *α* locus, as seen in Fig.S6.

4-way DMIs correspond to an interaction where all four loci require a particular combination of alleles for good patterning. As previously noted they are of much smaller number compared to 2- and 3-way DMIs, so here, we do not examine these DMIs in detail. However, we note that the 4-way DMIs shown in Fig.S7, show a similar pattern as found with 3-way DMIs, where for small scaled population sizes the DMIs tend to cluster, which suggests, as found for 2- and 3-way DMIs, this is due to sequence entropy constraints dominating the growth of DMIs; on the other hand, at large scaled population sizes this degeneracy is lifted and each hybrid has a different growth rate of DMIs, depending on their particular contribution to fitness and how that balances against the constraints of sequence entropy.

**FIG. S7.**
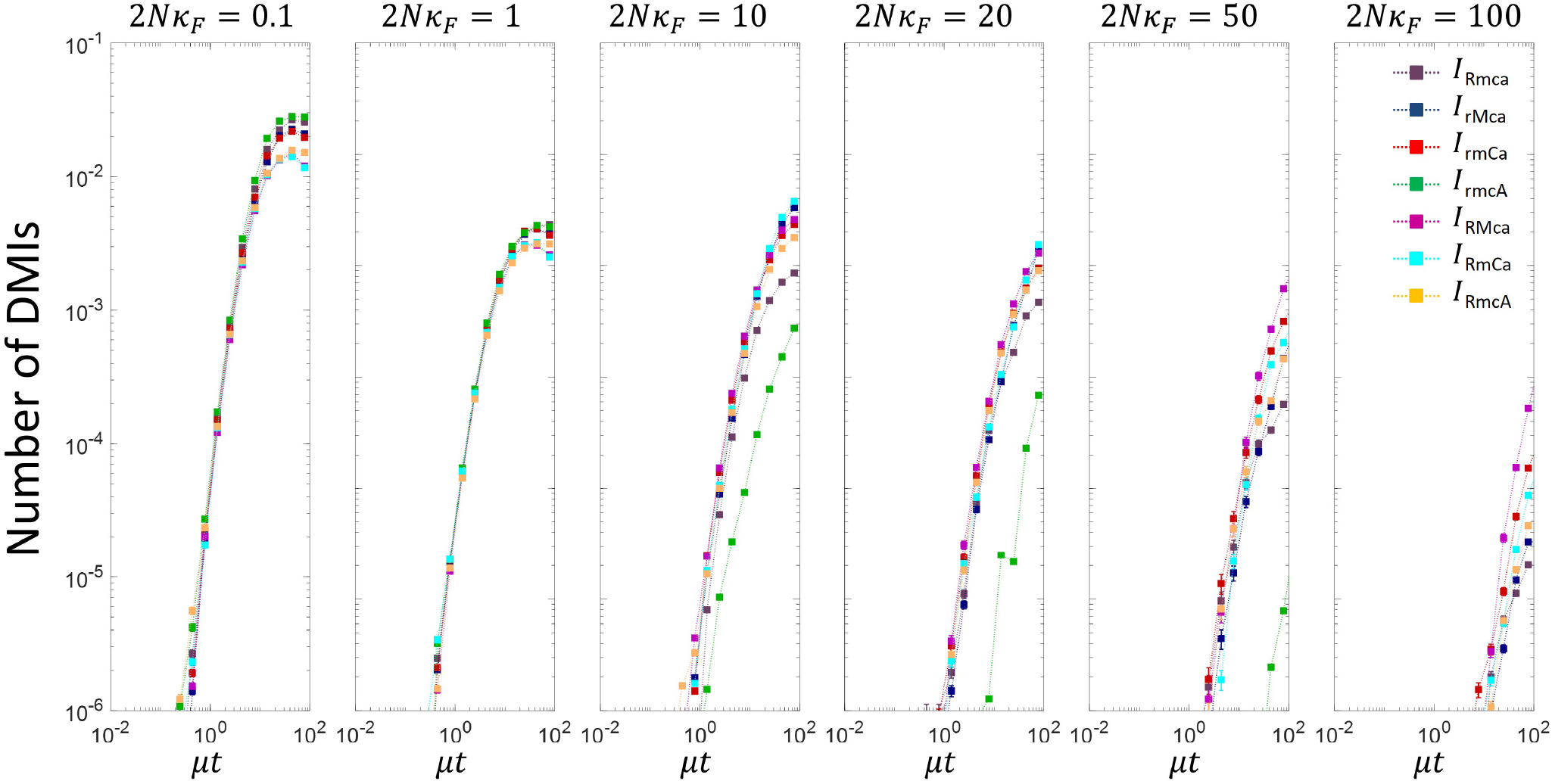
Plot of the spectrum 4-way DMIs vs divergence time for different scaled population sizes.

